# *Medicago truncatula* MOT1.3 is a plasma membrane molybdenum transporter required for nitrogenase activity in root nodules

**DOI:** 10.1101/102517

**Authors:** Manuel Tejada-Jiménez, Patricia Gil-Díez, Javier León-Mediavilla, Jiangqi Wen, Kirankumar S. Mysore, Juan Imperial, Manuel González-Guerrero

## Abstract

- Molybdenum, as a component of the iron-molybdenum cofactor of nitrogenase, is essential for symbiotic nitrogen fixation. This nutrient has to be provided by the host plant through molybdate transporters.
- Members of the molybdate transporters family MOT1 were identified in the model legume *Medicago truncatula* and their expression in nodules determined. Yeast toxicity assays, confocal microscopy, and phenotypical characterization of a *Tnt1* insertional mutant line were carried out in the one *M. truncatula* MOT1 family member expressed specifically in nodules.
- Among the five MOT1 members present in *M. truncatula* genome, *MtMOT1.3* is the only one uniquely expressed in nodules. MtMOT1.3 shows molybdate transport capabilities when expressed in yeast. Immunolocalization studies revealed that MtMOT1.3 is located in the plasma membrane of nodule cells. A *mot1.3-1* knockout mutant showed an impaired growth concomitant with a reduction in nitrogenase activity. This phenotype was rescued by increasing molybdate concentrations in the nutritive solution, or upon addition of an assimilable nitrogen source. Furthermore, *mot1.3-1* plants transformed with a functional copy of *MtMOT1.3* showed a wild type-like phenotype.
- These data are consistent with a model in which MtMOT1.3 would be responsible for introducing molybdate into nodule cells, which will be later used to synthesize functional nitrogenase.

## Introduction

Molybdenum is one of the scarcest oligonutrients in the biosphere (Esteifel, 2002). Plants use it as part of the molybdenum cofactor (Moco) in just five enzymes involved in nitrate assimilation, purine metabolism, phytohormone production, and sulfite detoxification (Tejada-Jimenez et al., 2013). This nutrient, unlike other transition metals, is recovered from soil as the oxyanion molybdate instead of a cationic form. This determines that the transporters involved in this process are not the classical ones required for iron, copper, or zinc uptake, but members of other families. Molybdate shares some physicochemical characteristics with sulfate leading to cross-inhibition of sulfate transport by molybdate, probably due to non-specific molybdate transport through sulfate transporters in plants (Stout et al., 1951). Therefore, until recently it was thought that sulfate transporters might mediate molybdate transport in eukaryotic systems (Mendel & Hansch, 2002; Kaiser et al., 2005). The only known plant-type specific molybdate transporters belong to the Molybdate Transporter type 1 (MOT1) family and were identified in parallel in the green alga *Chlamydomonas reinhardtii* (Tejada-Jimenez et al., 2007) and in the higher plant *Arabidopsis thaliana* (Tomatsu et al., 2007). They share a high degree of homology with the sulfate transport family SULTR, but lack the conserved STAS domain (Tejada-Jimenez et al., 2007). In *C. reinhardtii*, CrMOT1 is responsible for high-affinity molybdate uptake, a process that is not severely affected by sulfate, indicating that MOT1 proteins are molybdate-specific transporters (Tejada-Jimenez et al., 2007). In Arabidopsis two members of the MOT1 family have been identified. One of them has been proposed to play a role in efficient Mo uptake from the soil (Tomatsu et al., 2007), although this function is not clear given the conflicting subcellular localizations reported for this transporter in plasma membrane or in mitochondria (Tomatsu et al., 2007; Baxter et al., 2008); while a second member of the MOT1 family is located in the vacuole of leaves and seems to be involved in intracellular and inter-organ Mo transport (Gasber et al., 2011). More recently, a MOT1 protein (LjMOT1) has been identified in *Lotus japonicus*, with a role in molybdate uptake from soil and translocation to the shoots, being located in the plasma membrane when expressed in tobacco leaves (Gao et al., 2016). In addition, another molybdate transporter family, MOT2, belonging to the major facilitator superfamily has been identified in *C. reinhardtii* (Tejada-Jimenez et al., 2011), indicating that molybdate transporters have appeared at least twice in evolution. However, its functionality as molybdate transporters has only been proved in this alga.

While all the plants employ molybdenum for Moco biosynthesis, legumes have an additional use for it. These organisms also require molybdenum for the assembly of the iron-molybdenum cofactor (FeMoco) of nitrogenase (Georgiadis et al., 1992; Rubio & Ludden, 2008), the enzyme responsible for nitrogen fixation in their root nodules. In legumes, FeMoco is assembled by diazotrophic bacteria living within differentiated root organs, the nodules. These organs are developed in a complex process starting with the detection of rhizobial nodulation factors (Nod) by the host plant that leads to root hair curling, bacteria trapping, hydrolysis of the plant cell wall and bacteria delivery to the root nodule primordium through an infection thread (Kondorosi et al., 1984; Brewin, 1991; Oldroyd, 2013). Once in the plant cytoplasm, rhizobia together with a plant-derived membrane result in organelle-like structures, the symbiosomes, where nitrogen fixation takes place. Two different types of nodules can be found in legumes (Sprent, 2007): determinate and inderterminate nodules. In inderterminate nodules, as those found in *Medicago truncatula*, the continuous meristem growth results in the formation of at least four different zones in mature nodules: the meristematic zone that drives nodule growth; the infection/differentiation zone, where rhizobia are released through the infection thread and differentiates to bacteroids; the fixation zone where nitrogenase carries out its enzymatic activity; and the senescent zone where bacteroids are degraded (Vasse et al., 1990). An additional nodule zone, the interzone, can be defined as the transition between infection/differentiation and fixation zone (Roux et al., 2014). Once the oxygen tension drops in the interzone, endosymbiotic bacteroids express nitrogenase and the machinery that allows them synthesize FeMoco. Among them, is the *modABC* operon, responsible for molybdate uptake from the peribacteroid space (Maupin-Furlow et al., 1995; Delgado et al., 2006; Hernandez et al., 2009). Consequently, for molybdate to reach the bacteroids, it has to cross the plasma and the symbiosome membranes, a process that has to be mediated by transporters belonging to two different families, given the two different directions of transport required.

In spite of the essential role that molybdenum has in nitrogenase, and the importance of nitrogenase in legume colonization of new environments and in sustainable agriculture, the transporters involved in molybdenum supply to the nodule cells are not known. It can be hypothesized that molybdate transfer could in some cases be mediated by sulfate transporters. In this case, molybdate transfer across the symbiosome membrane could be carried out by SST1-like proteins, that have been previously associated with sulfate delivery to symbiosomes (Krusell et al., 2005). At the plasma membrane, another sulfate transporter, such a SHST1 homologue could mediate molybdate uptake by rhizobia-infected cells since this transporter from the legume *Stylosanthes hamata* has already been shown to transport molybdate when it is expressed in yeast (Fitzpatrick et al., 2008). However, *in planta* SHST1 is expressed in the root and it is involved in sulfate uptake (Smith et al., 1995), while its relationship with plant molybdate transport has not been determined yet. A more likely alternative would be transporters from the MOT1 or MOT2 families, since they could finely tune molybdate delivery for symbiotic nitrogen fixation, given their high specificity for this anion.

In this work, we have identified a *M. truncatula* member of the MOT1 family (MtMOT1.3) involved in molybdate transport to nodule cells. *MtMOT1.3* is specifically expressed in nodules. Its protein product is located in the plasma membrane of infected and non-infected cells in the fixation zone of the nodule, coinciding with the zone where *MtMOT1.3* is expressed. *M. truncatula* plants lacking a functional *MtMOT1.3* gene show a reduction of nitrogenase activity connecting MtMOT1.3 function with molybdenum supply for FeMoCo biosynthesis. This is the first molybdate transporter known to be specific of legume symbiotic nitrogen fixation.

## Materials and Methods

### Biological material and growth conditions

*M. truncatula* R108 seeds were scarified in the presence of concentrated sulfuric acid for 7 min. Then, seeds were washed several times with cold water, surface sterilized with 50 % (v/v) bleach for 90 s and incubated overnight with sterile water in the dark. After 48 h at 4 ºC, seeds were germinated in water-agar plates 0.8 % (w/v). Seedlings were transferred to sterile perlite pots or to Jenner’s solution for hydroponic growth (Brito et al., 1994), and inoculated with *Sinorhizobium meliloti* 2011 or the same bacterial strain transformed pHC60 (Cheng & Walker, 1998). Plants were grown in a greenhouse under 16 h light and 22 ºC. In the case of perlite pots, plants were watered every two days with Jenner’s solution or water alternatively. Nodules were collected 28 dpi. Plants growing under non-symbiotic conditions were supplemented every two weeks with 2 mM KNO_3_*. Agrobacterium rhizogenes* strain ARqua1 having the appropriate vector was used for *M. truncatula* hairy-root transformation (Boisson-Dernier et al., 2001). Agroinfiltration for transitory expression experiments were performed in *N. benthamiana* leaves using *A. tumefaciens* C58C1 carrying the corresponding genetic construct.

*Saccharomyces cerevisiae* strain 31019b (MATa *ura3 mep1Δ mep2Δ::LEU2 mep3Δ:: KanMX2*) was used for heterologous expression assays (Marini et al., 1997). Yeasts were grown in synthetic dextrose (SD) or yeast peptone dextrose medium supplemented with 2 % glucose (Sherman et al., 1986).

### Quantitative real-time RT-PCR

Transcriptional expression studies were carried out by real-time RT-PCR (StepOne plus, Applied Biosystems) using the Power SyBR Green master mix (Applied Biosystems). Primers used are indicated in Supporting Information, Table S1. RNA levels were normalized by using the *ubiquitin carboxy-terminal hydrolase* gene as internal standard. RNA isolation and cDNA synthesis were carried out as previously described (Tejada-Jimenez et al., 2015).

### GUS Staining

*pMtMOT1.3::GUS* construct was obtained by amplifying 1.1 kb upstream of the *MtMOT1.3* (Nakagawa et al., 2007) start codon using the primers 5MtMOT1.3pGW and 3MtMOT1.3pGW (Supporting Information, Table S1). The resulting fragment was cloned by Gateway cloning technology (Invitrogen) into pGWB3 vector. Hairy-root transformation was carried out as described above. GUS staining was performed in root of 28-dpi plants as previously described (Vernoud et al., 1999). Nodules sections were clarified with 50% bleach for 30 min.

### Immunohistochemistry and confocal microscopy

A genomic region comprising *MtMOT1.3* full gene and 1.1 kb upstream of its start codon was cloned into pGWB13 vector (Nakagawa et al., 2007) using Gateway cloning technology (Invitrogen). The resulting genetic construct contains *MtMOT1.3* gene under the control of its own promoter and with three C-terminal HA epitopes in frame. Hairy-root transformation was carried out as indicated above. *M. truncatula* transformed plants were inoculated with *S. meliloti* 2011 constitutively expressing GFP. Nodules and roots were collected at 28-dpi and fixed at 4 ºC overnight in 4 % (w/v) paraformaldehyde and 2.5 % (w/v) sucrose in phosphate-buffered saline (PBS). Fixed plant material was sectioned, 100 µm wide, with a Vibratome 1000 Plus. Sections were dehydrated by serial incubation with methanol (30 %, 50 %, 70 % and 100 % [v/v] in PBS) for 5 min and then rehydrated following the same methanol series in reverse order. Cell wall permeabilization was carried out by incubation with 2 % (w/v) cellulase in PBS for 1 h and 0.1 % (v/v) Tween 20 for 15 min. Sections were blocked with 5 % (w/v) bovine serum albumin in PBS and then incubated with 1:50 anti-HA mouse monoclonal antibody (Sigma) in PBS at room temperature for 2 h. Primary antibody was washed three times with PBS for 10 min and subsequently incubated with 1:40 Alexa594-conjugated anti-mouse rabbit monoclonal antibody (Sigma) in PBS at room temperature for 1 h. Secondary antibody was washed three times with PBS for 10 min, and then DNA was stained using DAPI. Images were obtained with a confocal laser-scanning microscope (Leica SP8).

### Transient expression in *Nicotiana benthamiana* leaves

MtMOT1.3 transient expression in tobacco leaves was carried out as previously described (Voinnet et al., 2003). *MtMOT1.3* CDS was cloned into pGWB5 (Nakagawa et al., 2007) by Gateway cloning technology (Invitrogen), resulting in C-terminal fusion to GFP. *A. tumefaciens* C58C1 (Deblaere et al., 1985) cells independently transformed with this construct, with the plasma membrane marker pm-CFP pBIN (Nelson et al., 2007) or with the silencing suppressor p19 of *Tomato bushy stunt virus* (Voinnet et al., 2003) were injected into 4-week-old *N. benthamiana* leaves. Expression of the appropriate construct was analyzed after 3 d by confocal laser-scanning microscopy (Leica).

### Nitrogenase activity

Acetylene reduction assay was used to measure nitrogenase activity (Hardy et al., 1968). Wild-type and mutant plants 28 dpi were separately introduced in 30-mL tubes. Each tube was sealed with rubber stoppers and contained at least four independently transformed plants. Three mililiters of air of each bottle was replaced by the same volume of acetylene, tubes were subsequently incubated for 30 min at room temperature. Produced ethylene was measured by analyzing 0.5 mL of gas from each bottle in a Shimadzu GC-8A gas chromatograph using a Porapak N column.

### Metal content determination

Inductively coupled plasma optical emission spectrometry was carried out at the Unit of Metal Analysis in the Scientific and Technology Centers of the Universidad de Barcelona (Spain).

### Nitrate reductase activity

Nitrate reductase activity was analyzed as described by Scheible et al. (1997) with some modifications. We started from 100 mg of fresh leaves. The extraction buffer used for crude extracts preparation contained 100 mM potassium phosphate pH 7.5, 5 mM magnesium acetate, 10 % glycerol (v/v), 10 % polyvinylpolypyrrolidone (w/v), 0.1 % Triton X-100, 1 mM EDTA, 0.05 % ß-mercaptoethanol, 1 mM PMSF. Plant material was homogenized with liquid nitrogen and 1:6 extraction buffer (v/v). The crude extract was centrifuged at 14,000 x*g* at 4 ºC for 15 min. The reaction was initiated by adding 0.05 mL of crude extract to 0.5 mL of reaction buffer and incubated at 30 ºC for 20 min. The reaction buffer contained 50 mM potassium phosphate pH 7.5, 10 mM KNO_3_, 5 mM EDTA and 0.5 mM NADH. Nitrate reduction reaction was stopped by adding 1 volume of 1 % sulfanilamide in 2.4 M HCl, and 1 volume of 0.02 % N-1-naphtyl-ethylenediamine. After centrifugation absorbance at 540 nm was measured in UV/visible spectrophotometer Ultrospect 3300 pro (Amersham Bioscience).

### Bioinformatics

Members of the MOT1 family in *M. truncatula* were identified by the Basic Local Alignment Search Tool in the *M. truncatula* Genome Project database (http://www.jcvi.org/medicago/index.php), using as reference sequence a member of the MOT1 family from Arabidopsis (NP_180139). Protein sequence alignment and unrooted tree visualization were performed with MEGA7 package (http://www.megasoftware.net/) using ClustalW software and neighbor joining algorithm. Accession numbers: *Arabidopsis thaliana* AtMOT1.1, NP_180139.1; AtMOT1.2, NP_178147.1; AtSULTR1.1, NP_192602.1; AtSULTR2.1, NP_196580.1; AtSULTR3.1, ANM64117.1; AtSULTR4.1, AED91910.1; *Brassica napus* BnMOT1.1, XP_013709878.1; BnMOT1.3, XP_013667816.1; BnSULTR1.1, XP_013715602.1; BnSULTR2.1, NP_001302517.1; BnSULTR3.1, XP_013737126.1; BnSULTR4.1, XP_013667920.1; *Glycine max* GmMOT1.1, XP_003545516.1; GmMOT1.4, KRH05029.1; GmMOT1.6, XP_003527804.1; GmMOT1.7, XP_003523708.1; GmSULTR1.3, XP_006593569.1; GmSULTR2.1, XP_003531538.1; GmSULTR3.1, XP_003521258.1; GmSULTR4.2, XP_003552670.1; *Lotus japonicus* LjMOT1.1, AFK43331.1; LjMOT1.2, AJE26312.1; LjSST1, CAL36108.1; *Medicago truncatula* MtMOT1.1, XP_013465770.1; MtMOT1.2, XP_013465776.1; MtMOT1.3, XP_013460259.1; MtMOT1.4, XP_013454709.1; MtMOT1.5, XP_003603486.1; MtSULTR1.1, XP_003614968.1; MtSAT1, XP_003602002.1; *Oryza sativa* OsMOT1.1, XP_015650610.1; OsMOT1.2, XP_015621613.1; OsSULTR1.2 XP_015650733.1; OsSULTR2.1, ABF94445.1; OsSULTR3.1, BAS82415.1; OsSULTR4.1, XP_015612472.1; *Phaseolus vulgaris* PvMOT1.1, XP_007161270.1; PvMOT1.2, XP_007137089.1; PvMOT1.3, XP_007161272.1; PvMOT1.4, XP_007135722.1; PvSULTR1.1, XP_007141140.1; PvSULTR2.1, XP_007163633.1; PvSULTR3.1, XP_007162459.1; PvSULTR4.1, XP_007139276.1; *Vitis vinifera* VvMOT1.1, XP_002281989.2; VvMOT1.2, XP_002285217.1; VvSULTR1.1, XP_010664070.1; VvSULTR2.1, XP_010652824.1; VvSULTR3.1, XP_003632327.1; VvSULTR4.2, XP_002282491.2.

### Statistical tests

Data were analyzed by Student’s unpaired t test to calculate statistical significance of observed differences. Test results with p-values less than 0.05 were considered as statistically significant.

## Results

### *MtMoT1.3* belongs to the molybdate transporter MOT1 family and is specifically expressed in nodules

Due to the possible role of MOT1 transporters in Mo supply for symbiotic nitrogen fixation, we carried out a search for MOT1 members in the *M. truncatula* genome. Five putative *M. truncatula* proteins were found (encoded by *Medtr1g010210*, *Medtr1g010270*, *Medtr3g464210*, *Medtr4g011600* and *Medtr3g108190*) showing a sequence identity between 51.4 % and 65.7 % with an Arabidopsis member of the MOT1 family. These proteins were annotated as sulfate transporter-like proteins; however, sequence comparison showed that they cluster with plant MOT1 proteins and are more distant to sulfate transporters (SULTR) (Fig. 1a). Additionally, these proteins also contain the two sequence motifs conserved in all MOT1 proteins (Tejada-Jimenez et al., 2007) (Fig. 1b). Therefore, we named these proteins MtMOT1.1 to MtMOT1.5, respectively. The number of MOT1 members present in *M. truncatula* contrasts with the situation reported for Arabidopsis where only two MOT1 transporters are present (Tomatsu et al., 2007; Baxter et al., 2008; Gasber et al., 2011). Analysis of the number of MOT1 members in already sequenced plants showed that these proteins are present, in average, in a higher number in legumes than in non-legumes plants (Supporting Information, Table S2). This finding suggests a particular role of MOT1 proteins in legumes, where an additional molybdenum sink is present: diazotrophic bacteria in root nodules.

**Figure 1.**
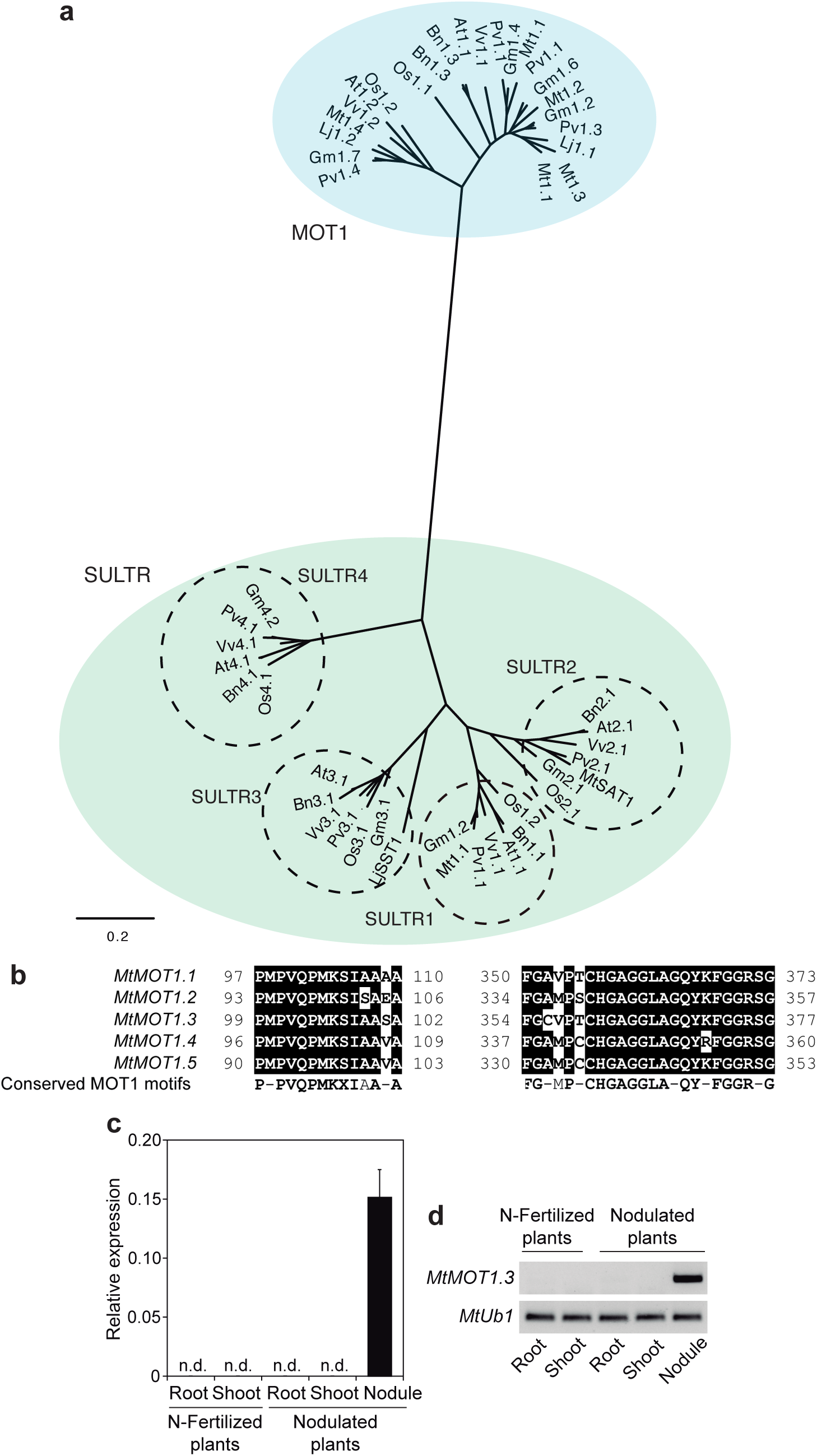
MtMOT1.3 is a member of the MOT1 protein and is expressed in the nodule. (a) Unrooted tree of the plant SULTR and MOT1 families. (b) Alignment of the conserved motifs of the MOT1 proteins present in all *M. truncatula* MOT1 members. Sequences were aligned using ClustalW method and it is extracted from a larger alignment including the full protein sequence of the *M. truncatula* MOT1 members. (c) Determination of *MtMOT1.3* expression in nodulated and nitrogen-fertilized *M. truncatula* plants relative to the internal standard gene Ubiquitin carboxyl-terminal hydrolase. Data are the mean ± SD of two independent experiments with 4 pooled plants. n.d., non-detected. (d) *MtMOT1.3* expression in nodulated and nitrogen-fertilized *M. truncatula* plants. *Ubiquitin carboxyl-terminal hydrolase1* (*MtUb1*) was used as control for PCR amplifications.

In order to study the possible relationship of the five putative MOT1 members of *M. truncatula* with symbiotic nitrogen fixation, a transcriptional analysis of these genes was carried out by real-time RT-PCR in plants during symbiotic association with *S. meliloti* or in uninoculated plants fertilized with nitrogen. We found that *MtMOT1.3* expression is restricted to the nodule (Fig. 1c and 1d), while transcripts from the other four *MtMOT1* genes were also detected at varying levels in other organs of the analyzed plants (Supporting Information, Fig. S1). These results strongly suggest an important role of MtMOT1.3 in the nodule, likely in molybdate transport connected to symbiotic nitrogen fixation. *MtMOT1.3* expression levels were not significantly affected by molybdate concentration in the nutritive solution (Supporting Information, Fig. S2).

### MtMOT1.3 is a molybdate transporter

All the MOT1 proteins reported so far mediate molybdate transport, in form of the oxyanion molybdate, to the cytosol (Tejada-Jimenez et al., 2013). The high sequence similarity that MtMOT1.3 shares with the already characterized molybdate transporters suggests that this protein could also mediate molybdate transport in the same direction. To test this hypothesis, *MtMOT1.3* was heterologously expressed in the yeast *S. cerevisiae*. This yeast is a good model to study of Mo transporters since it is one of the few organisms that do not use molybdenum, excluding any effect of endogenous specific molybdate transport activity (Mendel & Bittner, 2006). Toxicity studies showed that yeasts expressing *MtMOT1.3* exhibit a defective growth in synthetic dextrose (SD) solid medium in the presence of 50 µM molybdate, compared with yeast transformed with the empty vector pDR196, but no growth differences were observed with the control when no additional molybdate was added to the growth medium (Fig. 2a). Similar results were obtained in SD liquid medium by monitoring yeast growth along the time (Fig. 2b; Supporting Information, Fig. S3). The toxic effect observed in yeast is the likely result of molybdenum being transported and accumulated in the cell as a result of MtMOT1.3 activity (Fig. 2c). This result supports the functionality of MtMOT1.3 as molybdate transporter towards the cytosol.

**Figure 2.**
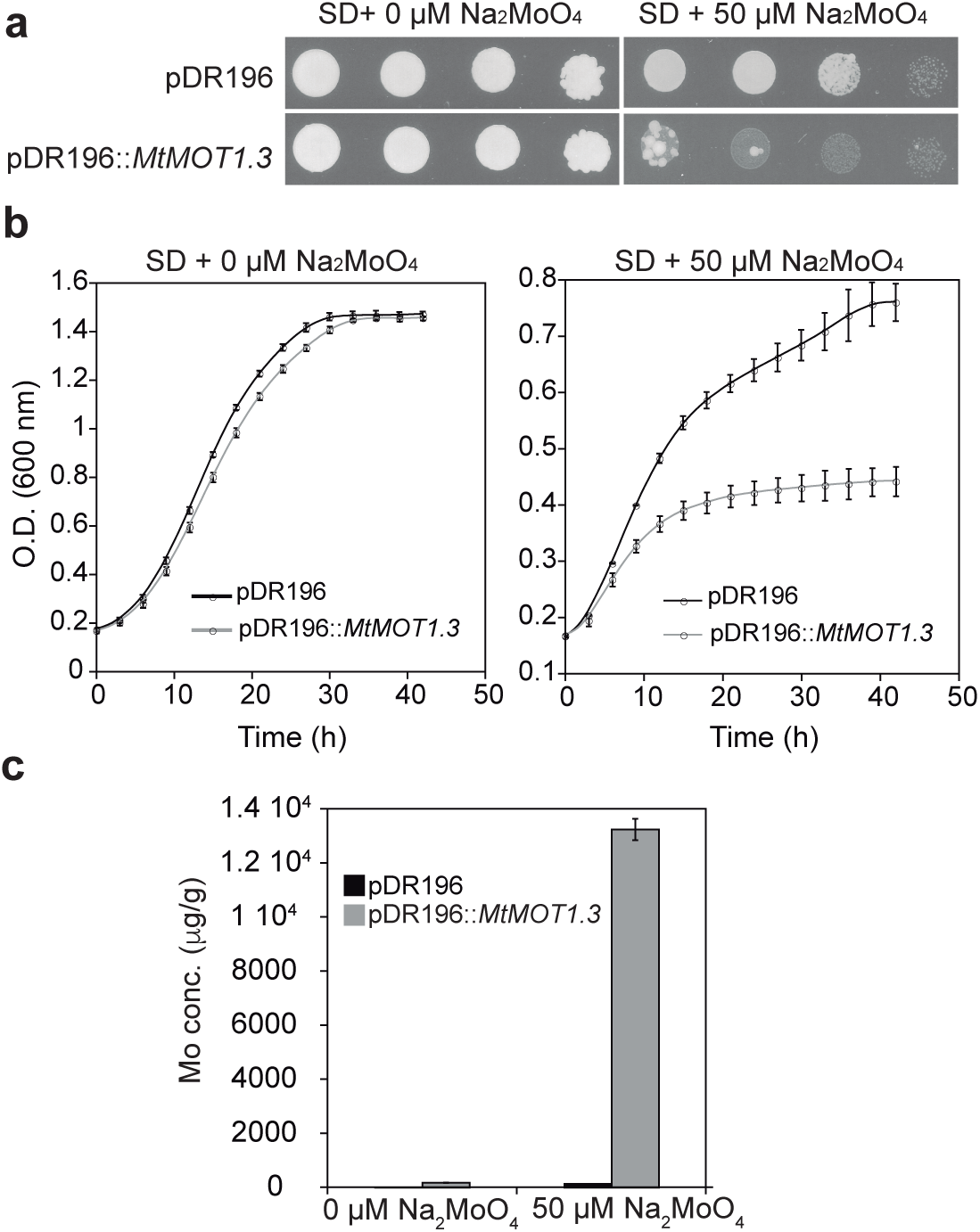
MtMOT1.3 transports molybdate towards the cytosol. (a) *S. cerevisiae* strain 31019b was transformed with PDR196 empty vector or PDR196 containing *MtMOT1.3* coding sequence, and grown at 28 ºC for 3d in serial dilution (10X) on SD solid medium medium without an added molybdenum source or containing 50 µM sodium molybdate. (b) *S. cerevisiae* strains used in (a) were grown in SD liquid medium, with and without 50µM sodium molybdate, at 28 ºC for 42 h. Yeast growth was monitored by measuring optical density at 600 nm every 3h. (c) Molybdenum content in the yeast grown in (b) after 42 h. Data are the mean ± SD of two independent experiments.

### *MtMOT1.3* is expressed in the plasma membrane of nodule cells of the interzone and fixation zones

To investigate the role of MtMOT1.3 in molybdenum supply to the nodule, we studied the tissue specific localization of *MtMOT1.3* expression. This was assessed by analyzing the expression of the ß-glucoronidase (*gus*) gene under the control of 1.1 kb of genomic DNA directly upstream of the *MtMOT1.3* starting codon, that was selected as promoter. Using this genetic construct, GUS activity was found in the nodule interzone, with the maximum in the area corresponding with the fixation zone (Fig. 3a and 3b). This expression distribution within the nodule matches with the transcription data available in the Symbimics database obtained by means of laser-capture microdissection coupled to RNA sequencing (Roux et al., 2014), where *MtMOT1.3* expression is mainly detected in the interzone and fixation zones (Fig. 3c).

**Figure 3.**
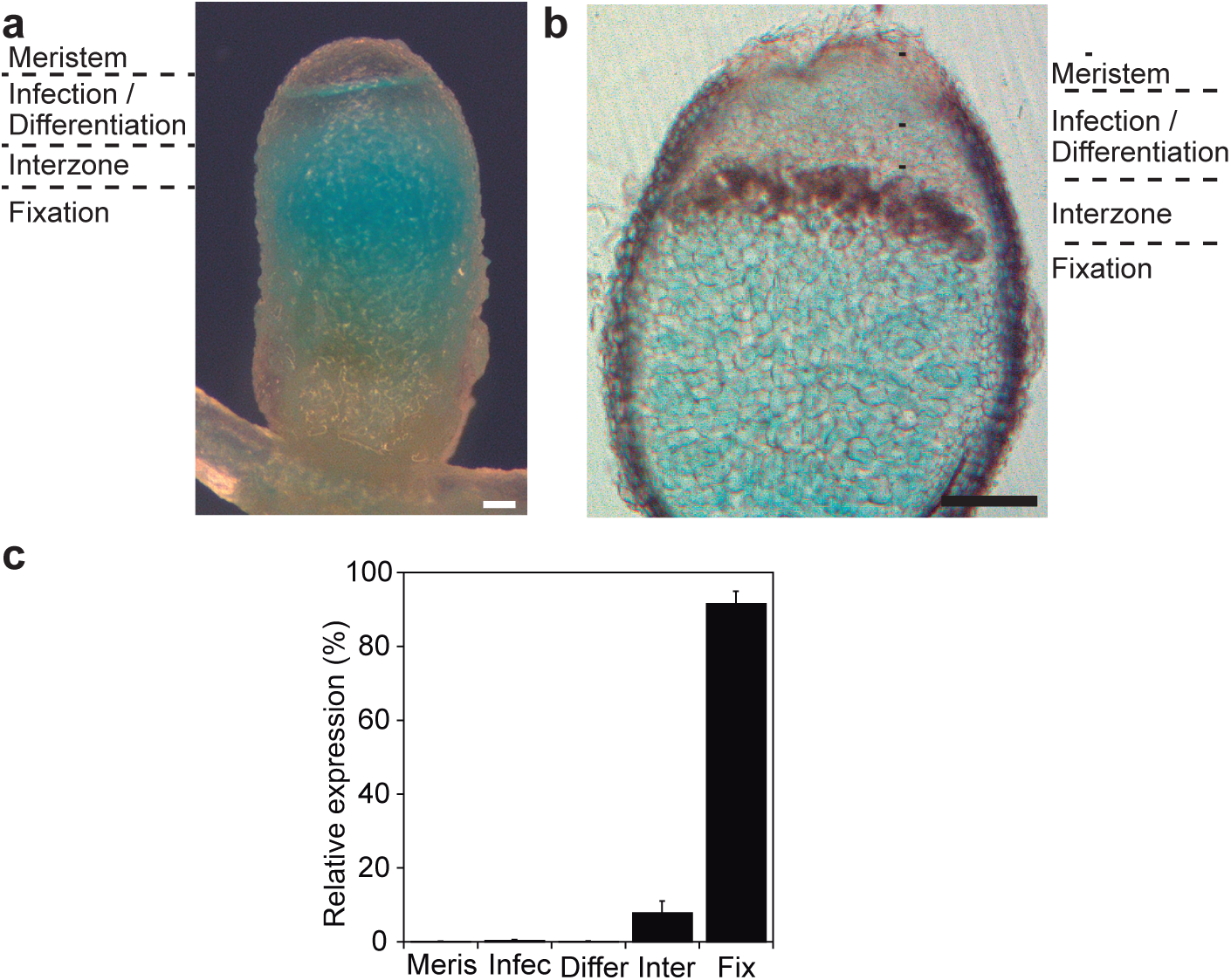
*MtMOT1.3* gene is expressed in the interzone and fixation zone of *M. truncatula* nodules. (a) and (b) GUS staining of *M. truncatula* 28-dpi nodules transiently expressing *gus* gene under the control of *MtMOT1.3* promoter. (a) Intact nodule. (b) Longitudinal nodule section. Bars = 200 µm. (c) Expression of *MtMOT1.3* in *M. truncatula* nodules by laser-capture microdissection coupled to RNA sequencing. Data were obtained from the Symbimics database (https://iant.toulouse.inra.fr/symbimics/). Meris, meristem; Infec, infection zone; Differ, differentiation zone; Inter, interzone; Fix, fixation zone.

Localization of MtMOT1.3 protein was also analyzed using an immunohistochemical and confocal microscopy approach. *M. truncatula* plants were transformed with a genetic construct comprising the genomic region of *MtMOT1.3* fused in frame with three hemagglutinin (HA) epitopes in C-terminus (MtMOT1.3-HA), under the control of the same promoter used for the GUS assay. The chimeric MtMOT1.3-HA protein was detected (in the red channel) with a mouse anti-HA antibody and an Alexa594-conjugated anti-mouse secondary antibody. Its position within the nodule was traced by DNA staining using 4’,6-diamino-phenylindole (DAPI) (blue) and a *S. meliloti* strain constitutively expressing GFP (green). In agreement with the GUS activity data, MtMOT1.3-HA was detected in the interzone and fixation zones of the nodule (Fig. 4a to 4c). Particularly, MtMOT1.3-HA signal was found in the periphery of infected and non-infected cells (Fig. 4d to 4f), fitting with plasma membrane localization. This signal was not the result of autofluorescence, since sections obtained from the same biological material, subjected to the same preparation protocol with the exception of the incubation with the primary anti-HA antibody, did not show any fluorescence in the Alexa594 emission spectrum (Supporting Information, Fig. S4). Moreover, the specific peripheral distribution of MtMOT1.3-HA was also observed using an Alexa488-conjugated secondary antibody in nodules containing m-Cherry-expressing *S. meliloti* (Supporting Information, Fig. S5). No MtMOT1.3-HA signal was detected by Western blot analyses in roots from nodulated plants (Supporting Information, Fig. S6), consistent with the nodule-specifc expression data obtained by real-time PCR. To confirm the subcellular localization of MtMOT1.3 in the plasma membrane, its coding sequence fused to the green fluorescent protein (GFP) in C-terminus was transiently expressed in *N. benthamiana* leaves together with a plasma membrane marker fused to cyan fluorescent protein (CFP). MtMOT1.3 signal and plasma membrane marker signal co-localized in *N. benthamiana* leaves cells expressing both genetic constructs (Fig. 4g to 4i), supporting the localization of MtMOT1.3 in the plasma membrane. However, no GFP or CFP signals were found in cells expressing only the plasma membrane marker or MtMOT1.3, respectively (Supporting Information, Fig. S7), thus ruling out any non-specific signal when both constructs are co-expressed.

**Figure 4.**
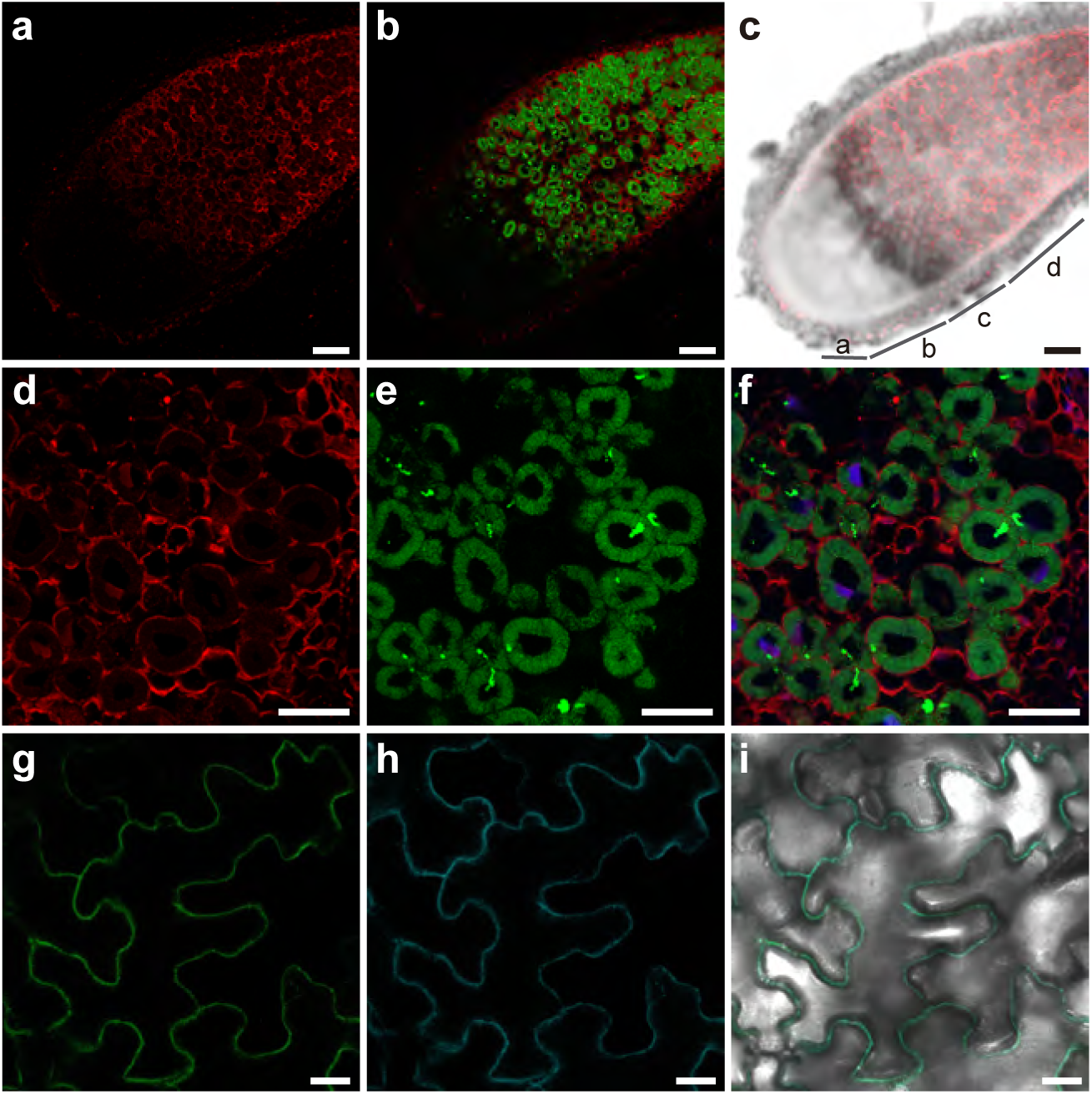
MtMOT1.3 is localized in the plasma membrane. (a) to (c) Cross section of a 28-dpi *M. truncatula* nodule transiently expressing MtMOT1.3-HA construct and infected with a *S. meliloti* strain constitutively expressing GFP (green). MtMOT1.3-HA was detected using an Alexa-594-conjugated antibody (red). (a) MtMOT1.3-HA signal. (b) Overlay of MtMOT1.3-HA and *S. meliloti* signals. (c) Overlay of MtMOT1.3-HA signal and differential interference contrast. a, meristem; b, infection/differentiation; c, interzone; b, fixation. Bars = 200 µm. (d) to (f) Closer view in the fixation zone of M. truncatula nodules. DNA was stained using DAPI (blue). (d) MtMOT1.3-HA signal. (e) *S. meliloti* signal. (f) Overlay of MtMOT1.3-HA, *S. meliloti* and DNA signals. Bars = 50 µm. (g) to (i) Transient co-expression of MtMOT1.3-GFP and plasma membrane marker-CFP in *N. benthamiana* leaves. (g) MtMOT1.3-GFP signal. (h) Plasma membrane marker-CFP signal. (i) Overlay of MtMOT1.3-GFP, plasma membrane marker-CFP signals and differential interference contrast. Bars = 25 µm.

### Lack of *MtMOT1.3* leads to a reduction of nitrogenase activity

In order to investigate the role of MtMOT1.3 in symbiotic nitrogen fixation, the mutant line NF10801 (*mot1.3-1*) was identified by a reverse genetics screening (Cheng et al., 2011; Cheng et al., 2014) from a *Transposable Element from N. tabacum* (*Tnt1*) insertion mutant library (Tadege et al., 2008). The mutant *mot1.3-1* carries a *Tnt1* insertion in the second exon of the *MtMOT1.3* gene at position +1782 from the starting codon (Fig. 5a). *MtMOT1.3* transcripts were not detected in the homozygous *mot1.3-1* mutant (Fig. 5b), therefore MtMOT1.3 activity is not present in this mutant line.

**Figure 5.**
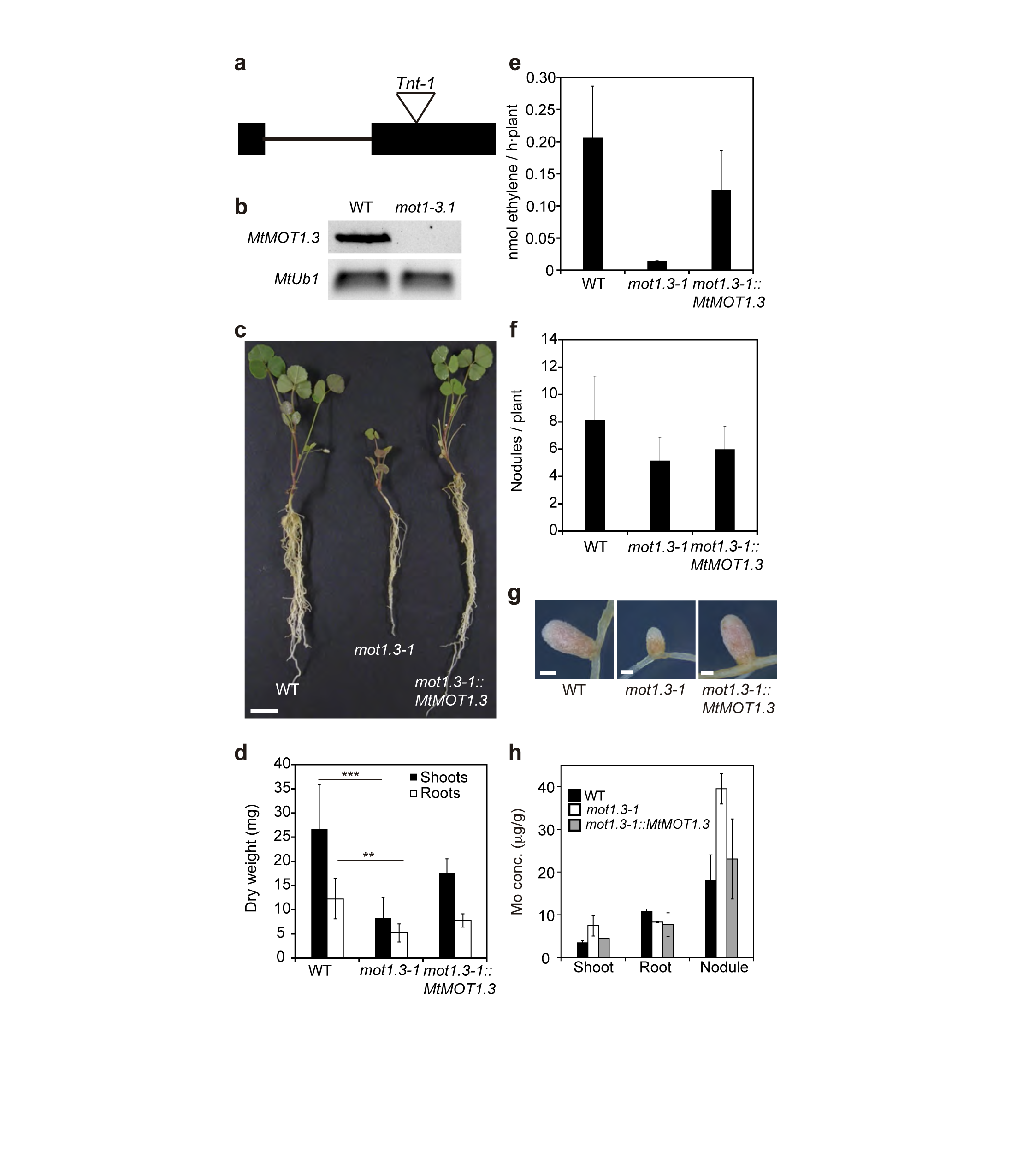
*MtMOT1.3* mutation results in a reduced nitrogen fixation rate. (a) Position of the *Tnt1* insertion within the *MtMOT1.3* genomic region. (b) RT-PCR amplification of *MtMOT1.3* transcript in 28 dpi nodules of *M. truncatula* wild type (WT) and mutant (*mot1.3-1*) plants. *Ubiquitin carboxyl-terminal hydrolase1* (*MtUb1*) was used as control for PCR amplifications. (c) Growth of representative plants of wild type, *mot1.3-1* mutant and *mot1.3-1* mutant transformed with the *MtMOT13-HA* construct. Bar = 3 cm. (d) Dry weight of shoots and roots. Data are the mean ± SD of at least 6 independently transformed plants. Asterisk indicates significant differences: **p < 0.01, ***p < 0.005. (e) Nitrogenase activity in 28-dpi nodules. Acetylene reduction was measured in duplicate from two sets of four pooled plants. Data are the mean ± SD. (f) Number of nodules per plant. Data are the mean ± SD of at least 6 independently transformed plants. (g) Representative nodules of each *M. truncatula* line. Bar = 500 μm. (h) Molybdenum content in shoots, roots, and nodules of wild type, *mot1.3-1* mutant, and *mot1.3-1* mutant transformed with the *MtMOT13-HA* construct. Data are the mean ± SD of 3 sets of 4 pooled plants.

The phenotype of *mot1.3-1* was evaluated in symbiotic conditions, with nitrogen fixation as the sole nitrogen source, and under low Mo availability. In these conditions, *mot1.3-1* mutant showed a reduced growth rate compared with wild-type plants (Fig. 5c). Consequently, plant biomass was reduced in the *mot1.3-1* mutant by 70 % in shoots and 55 % in roots (Fig. 5d). Nitrogenase activity was measured in *mot1.3-1* and wild-type plants by the acetylene-reduction assay (Dilworth, 1966; Schöllhorn & Burris, 1966). *mot1.3-1* plants showed a reduction of 90 % in nitrogenase activity, as compared to the activity measured in wild-type plants (Fig. 5e). On a per-nodule basis, *mot1.3-1* plants had in average 24 % of the nitrogenase activity of the wild-type ones (Supporting Information, Fig. S8). Nodulation was not affected in the *mot1.3-1* mutant, since nodules were comparable to wild-type plants in terms of number and shape (Fig. 5f and 5g). Moreover, no differences in nodule anatomy, or nodulation kinetics were observed between mutant and control plants (Supporting Information, Fig. S9). However, these nodules were on average smaller than those from wild type plants (Fig. 5g; Supporting Information, Fig. S10). In addition, nodule molybdenum content in *mot1.3-1* plants was significantly higher than the control (Fig. 5h), indicative of a role on molybdenum homeostasis in this organ. In order to investigate whether the phenotype of *mot1.3-1* plants is caused by a shortage in molybdenum supply, these plants were watered with a nutrient solution containing 5 µg/L ammonium heptamolybdate. Under molybdenum sufficient conditions, *mot1.3-1* mutants exhibited a growth rate, biomass and nitrogenase activity comparable to wild-type plants (Supporting Information, Fig. S10). A similar result was obtained in hydroponics, a growth condition where molybdenum concentrations can be better controlled (Supporting Information, Fig. S11). Interestingly, no MOT1 family member was more highly expressed in *mot1.3-1* nodules than in wild-type plants (Supporting Information, Fig. S12).

To validate that the phenotype observed was the result of the *Tnt1* insertion in *MtMOT1.3, mot1.3-1* plants were transformed with *MtMOT1.3-HA* construct under its own promoter (same genetic construct used for MtMOT1.3 immunolocalization assay). Insertion of the mutated gene in the *mot1.3-1* mutant under low molybdenum conditions also restored wild-type growth (Fig. 5a to 5e), supporting that mutation of *MtMOT1.3* is responsible for this abnormal phenotype. In addition, it also validates the immunofluorescence data, since without a wild type-like expression profile and localization, no complementation would have been observed.

To check whether the lack of MtMOT1.3 activity could affect other processes in the plant, the phenotype of *mot1.3-1* mutant was assayed under non-symbiotic conditions, with nitrate as nitrogen source. In this situation, plant growth relies on the activity of the molybdo-enzyme nitrate reductase to reduce nitrate to nitrite that will be subsequently converted to ammonia and incorporated to amino acids by the glutamine synthetase/glutamine synthase pathway (Bernard & Habash, 2009). In these conditions *mot1.3-1* plants showed a phenotype similar to wild-type plants in terms of growth rate, biomass and nitrate reductase activity, regardless of the Mo availability (Fig. 6). Therefore, the function of MtMOT1.3 seems to be restricted to symbiotic conditions when symbiotic nitrogen fixation has an important role in plant growth.

**Figure 6.**
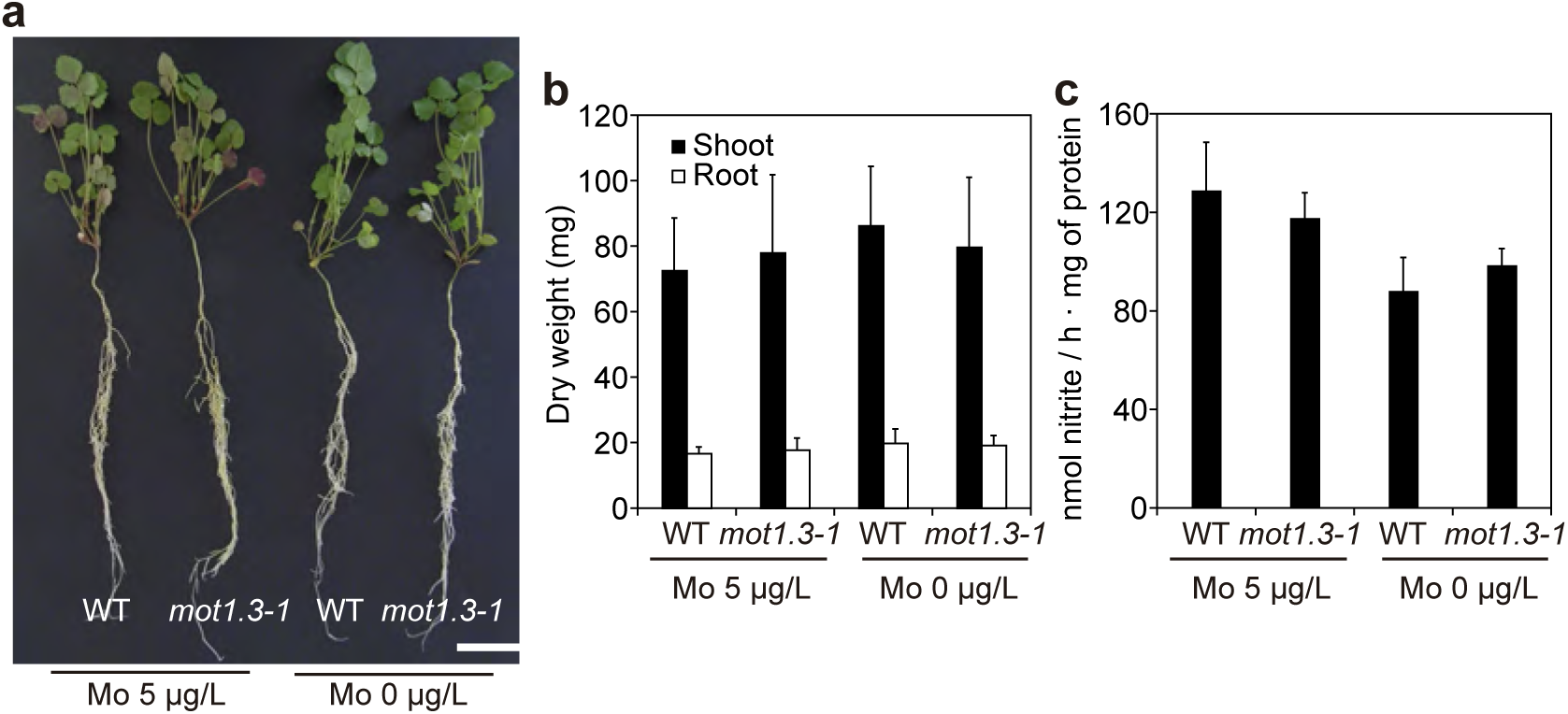
*mot1.3-1* mutant phenotype is not present in non-symbiotic conditions. (a) Growth of representative plants of wild type and *mot1.3-1* mutant. Bar = 3 cm. (b) Dry weight of shoots and roots. Data are the mean ± SD of at least 8 independent plants. (c) Nitrate reductase activity. Nitrate reduction was measured in duplicate. Data are the mean ± SD.

## Discussion

Obtaining nitrogen from the atmosphere by biological nitrogen fixation is a sustainable alternative to the intensive use of synthetic nitrogen fertilizers in agriculture (Smil, 1999). Legumes are able to use atmospheric N_2_ as nitrogen source by means of a symbiotic association with nitrogen-fixing bacteria from the soil. They occupy 12-15% of the world arable land (Graham & Vance, 2003). However, maintaining an adequate nitrogen fixation rate involves a complex biological process that requires an efficient nutrient supply from the plant to the rhizobia (Udvardi & Poole, 2013). Molybdenum supply is critical for symbiotic nitrogen fixation since this oligonutrient is needed to synthesize the enzyme nitrogenase, directly involved in the reduction of N_2_ to produce NH_4_^+^. In plants molybdenum is also present in other enzymes carrying out important metabolic processes such as nitrate assimilation (nitrate reductase) or phytohormone biosynthesis (aldehyde oxidase) (Tejada-Jimenez et al., 2013). Little is known about how plants take up molybdate from the soil and redistribute it to the sink organs, although members of the molybdate transport protein family MOT1 are very likely involved in this process in Arabidopsis (Tomatsu et al., 2007; Baxter et al., 2008; Gasber et al., 2011). However, their physiological role in molybdenum homeostasis is still unclear. Less is known for molybdate nutrition in legumes, in spite of the importance of molybdenum in symbiotic nitrogen fixation. Considering the renewed interest and recent advances towards introducing nitrogen fixing abilities into non-legumes (Charpentier & Oldroyd, 2010; Curatti & Rubio, 2014; Lopez-Torrejon et al., 2016) and the need for a more sustainable agriculture (Foley et al., 2011), this gap of our knowledge needs to be filled if we are to ensure proper delivery of molybdenum to produce functional nitrogenase in new biological systems. The present work represents a first but decisive step toward optimization of molybdate allocation for nitrogen fixation, where we have identified a nodule-specific MOT1 member in *M. truncatula* (MtMOT1.3) as responsible for molybdenum supply for symbiotic nitrogen fixation.

Legumes appear to have expanded the number of genes encoding MOT1 proteins in their genome in order to adapt to the presence of an additional molybdenum sink, the root nodule, doubling the average copy number in monocots and other dicots. This is in contrast to genes encoding Nramp transporters, some of which play an essential role in iron transport in rhizobia-infected cells (Kaiser et al., 2003; Tejada-Jimenez et al., 2015), where no significant increase in gene copy numbers were observed. One possible explanation for this, based on the fact that few plant proteins use molybdenum-based cofactors, is that MOT1 transporters are only expressed in very few cell types, highly specialized in the physiological processes catalyzed by Moco or FeMoco-dependent enzymes. If this hypothesis is right, nodule cells should express a specific MOT1 transporter responsible for ensuring molybdenum supply for symbiotic nitrogen fixation, as is the case for *MtMOT1.3* in *M. truncatula*.

*MtMOT1.3* is expressed in the nodule interzone and in the early fixation zone. This transcription pattern is also validated by the transcriptomic data obtained from the Symbimics database (Roux et al., 2014). This expression profile is consistent with a situation in which the rhizobia-infected cell is increasing its molybdenum content to transfer it to the bacteroids, so that when the physiological conditions are right, they can start synthesizing FeMoco. In contrast, iron-transporting *MtNramp1*, responsible for iron uptake by rhizobia-infected cells, shows a different expression profile, with a maximum of expression in the infection/differentiation zone (Tejada-Jimenez et al., 2015). This difference for the expression of iron and molybdenum transporter genes indicates that the uptake of these two elements occurs in two separate moments during nodule development, probably reflecting an earlier need for certain ferroproteins other than nitrogenase. It also indicates that molybdate release from the vasculature for uptake by rhizobia-infected cells occurs at the interzone and early fixation zone, in contrast to what has been proposed for iron and other transition metals (Rodriguez-Haas et al., 2013; González-Guerrero et al., 2014; González-Guerrero et al., 2016). However, neither *MtMOT1.3* nor *MtNramp1* are expressed in the older parts of the fixation zone. This suggests that no additional metal uptake is taking place in this zone, indicating either a high degree of protein stability in the fixation zone, or an effective recycling of essential metallic nutrients.

MtMOT1.3 is a plasma membrane-bound protein, as indicated by both the immunolocalization of a HA-tagged fusion in *M. truncatula* nodules and the colocalization studies of MtMOT1-GFP with a plasma membrane marker in tobacco leaves. In Arabidopsis, while AtMOT1.2 is clearly localized in the vacuole of leaf cells (Gasber et al., 2011), contradicting localization data have been reported for AtMOT1.1, claiming either plasma membrane or mitochondrial localizations (Tomatsu et al., 2007; Baxter et al., 2008). These conflicting data seem to be caused by the GFP marker fused to AtMOT1.1, since when it is fused to the N-terminus of AtMOT1.1 it leads to plasma membrane localization, while the C-terminus fusion results in mitochondrial localization. Our MtMOT1.3 subcellular localization data have been obtained using two different methodologies: immunolocalization in *M. truncatula* cells by detecting HA epitope fused to MtMOT1.3, and transient expression in *N. benthamiana* fused to GFP. Both approaches yielded the same result reinforcing MtMOT1.3 localization in the plasma membrane. The observed immunofluorescence data are not the result of any autofluorescence, since similar fluorescence profiles were observed using two different fluorescent probes. Moreover, when no probe was added to the sections, no fluorescence was detected in them, indicating that the signal was the result of the antibody binding. There is no evidence of unspecific recognition of any epitope other than the HA-tag in the Western blot analysis carried out using an anti-HA antibody. Furthermore, previous data showed that the Alexa594-conjugated antibody used was specific for HA-expressing proteins (Tejada-Jiménez et al., 2015). Finally, expressing *MtMOT1.3-HA* in a *mot1.3-1* mutant background resulted in complementation and recovery of the wild-type phenotype, which indicates that the tagged transporter is being targeted to the proper subcellular compartment.

Like MOT1 family members, MtMOT1.3 transports molybdate towards the cytosol, as indicated by the yeast toxicity assays. The phenotype characterization of a *Tnt1* mutant line strongly suggests that this activity is used by *M. truncatula* to introduce molybdate into nodule cells. The knockout mutant line *mot1.3-1* has lower nitrogenase activity than wild-type control plants, both when whole plants or nodules were compared. Consequently, plant growth is reduced due to lack of fixed nitrogen. Nodules from *mot1.3-1* are on average smaller than those from wild-type plants. This is not caused by a delayed nodulation, since nodulation kinetics are very similar to wild-type plants. Neither it is due to a defect of the nodulation process, as the nodule organization does not show any alteration in *mot1.3-1* nodules. Finally, the observed reduction in nodule size (around 75% of wild type) cannot account for the observed reduction in nodule nitrogenase activity (ca 25% of wild-type leves). Alternatively, it could be hypothesized that the reduction in nodule size could be caused by the reduced uptake of a nutrient essential for nodule functioning. The *mot1.3-1* phenotype is molybdenum-dependent, since molybdate fortification of the nutritive solution resulted in wild-type looking *mot1.3-1* plants, in apparent contradiction to the higher levels of molybdenum detected in *mot1.3-1* nodules. This accumulation pattern has also been observed when studying a nodule-specific plasma membrane copper transporter (unpublished data). One possible explanation is the existence of a signal indicating intracellular metal deficiency that would trigger more metal being transported to the nodule. Since the uptake transporter is not present, this would result in metal accumulation in the nodule apoplast and in the vasculature. In any case, the molybdate transport capabilities shown in yeast, the alteration of molybdate homeostasis in nodules, the reduction of a molybdate-dependent enzymatic activity in nodules, its plasma membrane localization, and the restoration of a wild-type phenotype by increasing molybdate concentrations in the nutritive solution, point out to a role of MtMOT1.3 in molybdate uptake by nodule cells.

The phenotype reversal by increasing the molybdate content of the nutritive solution is the result of other membrane transporters that at higher molybdate concentrations could counterbalance the absence of MtMOT1.3 activity. In *M. truncatula* all the MOT1 proteins are expressed within the nodule, although only *MtMOT1.3* exhibited a nodule-specific transcription pattern. It could be argued that another member of the MOT1 family could carry out this role. If this were the case, an induction of its expression levels in *mot1.3-1* nodules would be expected. However, we observe a slight reduction of their expression levels, suggesting a different role in molybdenum homeostasis in nodules. Alternatively, sulfate transporters could also compensate the lack of MtMOT1.3; for instance, the sulfate transporter SHST1 from the legume *Stylosanthes hamata* is able to transport molybdate upon expression in *S. cerevisiae* (Fitzpatrick et al., 2008). Furthermore, proteins of the MOT2 family have been also related to molybdate transport in the green alga *C. reinhardtii* (Tejada-Jimenez et al., 2011) and members of the MOT2 family are present in *M. truncatula* genome. However according to the database Symbimics, none of SHST1 or MOT2 *M. truncatula* orthologues are significantly expressed in nodules (Roux et al., 2014), which indicates that either another MOT1 transporter carried out this role in spite of the reduced expression levels, or a member of a yet-to-be-identified molybdate transporter family would be supplementing MtMOT1.3 function.

Under non-symbiotic conditions, MtMOT1.3 does not play an essential role, as indicated by its expression profile and by the lack of phenotype observed under these conditions. Even when the plants require Moco-dependent nitrate reductase activity to grow, no differences were observed between mutant and wild-type plants. These results reinforce the hypothesis of specialization of a legume MOT1 transporter during evolution to provide molybdenum for symbiotic nitrogen fixation.

In conclusion, molybdate transported by the host plant is released from the vasculature into the apoplast of the interzone/early fixation zone (Fig. 7). From there, MtMOT1.3 introduces molybdate into rhizobia-infected cells, as well as in the non-infected ones within the nodule. The latter raises the question of why the non-infected cells would require molybdenum. Possible explanations would be that they are being prepared if they become infected, that they are buffering or storing molybdenum to be used later, or that this molybdenum is required to synthesize active Moco-dependent enzymes in these cells. Once in the cell cytosol, molybdate has to be transported across the symbiosome membrane. Since no molybdate-specific efflux system is known, it could be speculated that this role is carried out by a sulfate transporter specific of this membrane such as SST1, whose mutation has a severe impact on nitrogenase activity (Krusell et al., 2005). Once in the peribacteroid space, ModABC would introduce molybdate into the bacteroid (Delgado et al., 2006; Cheng et al., 2016).

**Figure 7.**
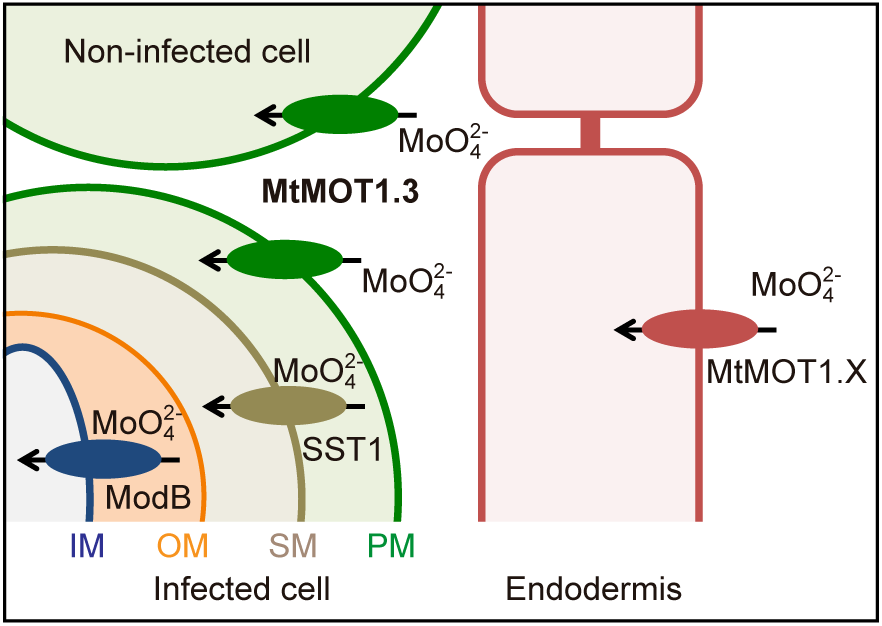
Proposed model for MtMOT1.3 function in molybdenum homeostasis in *M. truncatula*. Molybdenum, in form of molybdate would be released in the infection/differentiation zone, as already described for iron, in a process that could be assisted by a yet-to-be identified transporter that might belong to the MOT1 family (MtMOT1.X). From this point MtMOT1.3 would introduce molybdate into the cytoplasm of infected and non-infected cells within the nodule. The molybdate transport across the symbiosome membrane could be mediated by an orthologue of *L. japonicus* sulfate transporter SST1. Finally, molybdate would entry the bacteroids through the membrane component of the bacterial high affinity molybdate transport system ModB. IM, bacteroid inner membrane; OM, bacteroid outer membrane; SM, symbiosome membrane; PM, nodule cell plasma membrane.

## Acknowledgments

This research was funded by a European Research Council Starting Grant (ERC-2013-StG-335284), to MGG. Development of *M. truncatula Tnt1* mutant population was, in part, funded by the National Science Foundation, USA (DBI-0703285) to KSM. We would like to thank Dr. T. Coque and Dr. A. SanMillán (Ramón y Cajal Health Research Institute) for allowing us to use their Bioscreen equipment; and Dr. J. M. Argüello for sending us the mCherry-expressing *S. meliloti*. We would also like to acknowledge the other members of Laboratory 281 at Centro de Biotecnología y Genómica de Plantas (UPM-INIA) for their support and feedback in preparing this manuscript.

### Author contribution

MTJ carried out most of the experiments. PGD performed the confocal microscopy using the anti-HA Alexa488-conjugated antibody, the nodule development cross-section, and the molybdenum content in nodules. JLM carried out the expression of MOT1 genes in the *mot1.3-1* mutant, as well as studied the effecto of molybdate concentrations in transcription levels. Both PGD and JLM studied the nodulation process in wild type and *mot1.3-1* plants over time. JW and KSM performed *Medicago truncatula* mutant screening and isolated the *mot1.3-1* allele. MTJ, JI, and MGG designed the experiments, analyzed the data and wrote the article.

## Supplemental Methods

### Nodule development studies

Sterile seedlings were transferred to plates with Fahreus agar medium (Sauviac et al., 2005) without added molybdate and inoculated with 100 μl of saturated culture of *S. meliloti.* These plants were grown for 28 days at 16 h light and 22 ºC and 8 h darkness and 16 ºC. Morphological analyses were carried out in 28 dpi nodules collected and fixed in 0.25 % glutaraldehyde, 4 % formaldehyde, 2.5 % sucrose in 50 mM potassium phosphate buffer (pH7.4) at 4 ºC. Samples were dehydrated in an ethanol series and embedded in LR-White resin (London Resin Company Ltd, UK). Finally, nodules were placed in gelatin capsules, filled with resin and polymerized at 60 ºC for 24 h. Serial thin sections (0.5 μm) were cut with a Reichert Ultracut microtome (Leica, Vienna, Austria) fitted with a glass knife at the National Center for Electron Microscopy (Universidad Complutense de Madrid, Spain). Sections were stained with 1 % toluidine blue. Direct observation of sections was performed under a Zeis Axiophot photomicroscope (Carl Zeiss, Oberkoche, Germany) with an attached digital camera (Leica DFC 420C, Heerburgg, Switzerland).

### Immunohistochemistry and confocal microscopy

The localization of MtMOT1.3-HA was also determined using an anti-mouse Alexa488-conjugated antibody in sections from nodules colonized by a *S. meliloti* strain that constitutively expresses m-Cherry. Nodules and roots were collected at 28-dpi and fixed at 4 ºC overnight in 4 % (w/v) paraformaldehyde and 2.5 % (w/v) sucrose in phosphate-buffered saline (PBS). Fixed plant material was sectioned, 100 µm wide, with a Vibratome 1000 Plus. Cell wall permeabilization was carried out by incubation with 2 % (w/v) cellulase in PBS for 1 h and 0.1 % (v/v) Tween 20 for 15 min. Sections were blocked with 5 % (w/v) bovine serum albumin in PBS and then incubated with 1:50 anti-HA mouse monoclonal antibody (Sigma) in PBS at room temperature for 2 h. Primary antibody was washed three times with PBS for 10 min and subsequently incubated with 1:40 Alexa4884-conjugated anti-mouse rabbit monoclonal antibody (Sigma) in PBS at room temperature for 1 h. Secondary antibody was washed three times with PBS for 10 min, and then DNA was stained using DAPI. Images were obtained with a confocal laser-scanning microscope (Leica SP8).

### Western blot

Total root and nodule protein extracts were obtained from *M. truncatula* wild-type and from transformants expressing *MtMOT1.3-HA* as described by Larrainzar et al. (Larrainzar et al., 2007). Briefly, 100 mg fresh weight of nodules or roots was homogeneized in a mortar with liquid nitrogen and resuspended in cold extraction buffer (25 mM MES, 450 mM mannitol, 7 mM EDTA, 7 mM CaCl_2_, 5 mM MgCl_2_, 20 mM ascorbic acid, 10 mM ditithreitol pH7.2). Homogenates were briefly centrifuged at 500 x g for 5 min to remove cell debris. Protein determination was performed in accordance to Bradford (Bradford, 1976). 25 μg of this crude extract was loaded in SDS-PAGE gels (Laemmli, 1970), and protein bands visualized with Coomassie Brilliant Blue. HA-labelled proteins were detected by electroblotting the gels onto nitrocellulose membranes and immunostaining with a primary anti-HA mouse antibody (1:1000, Sigma) and a horseradish peroxidase-conjugated anti HA rabbit antibody (1:1000, Agrisera).

---

**Table S1.**
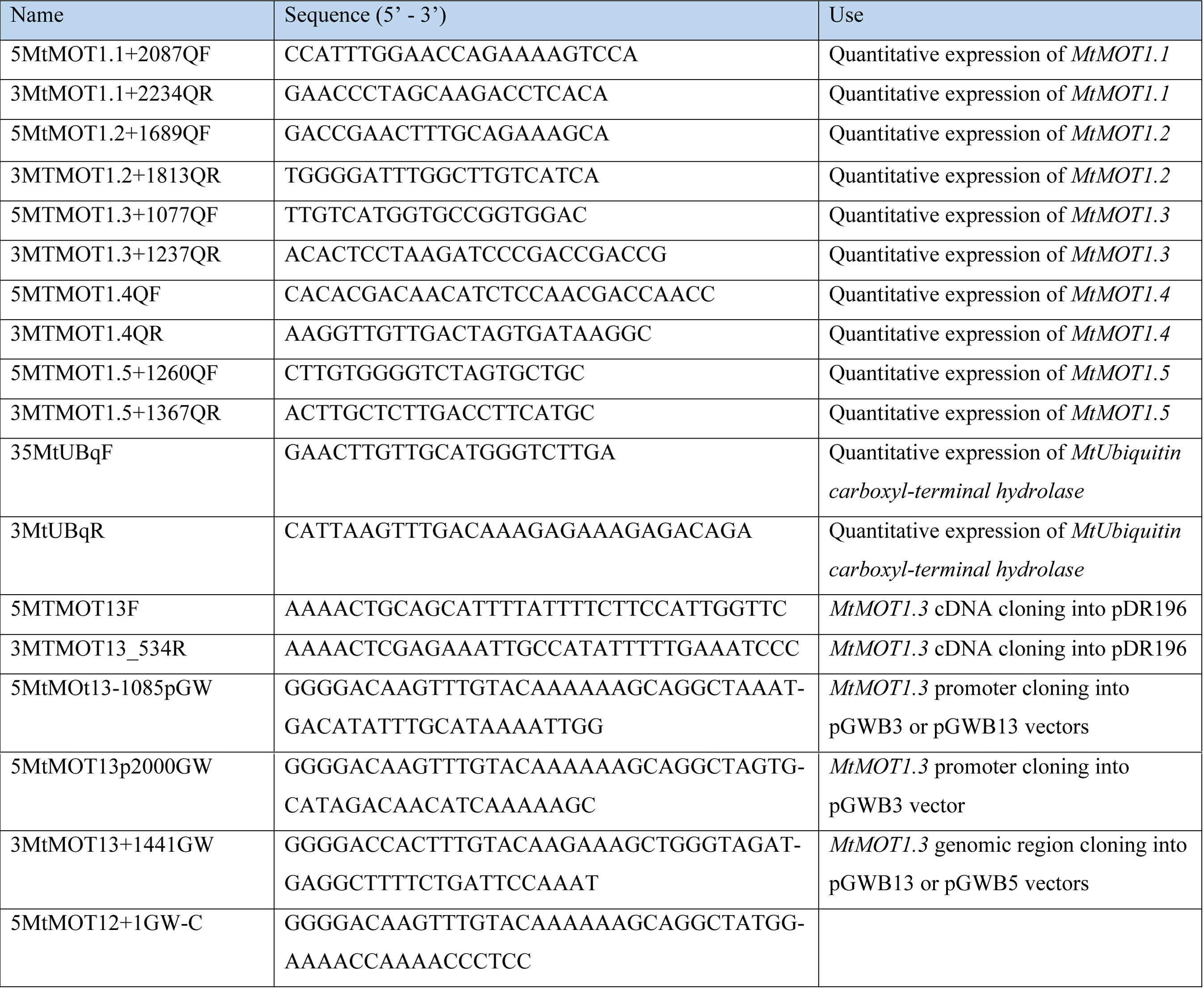
Primers used in this study

**Table S2.** Occurrence of MOT1 members in the genome of legume and non-legume plants.

| Legumes | Members of the MOT1 family |
| --- | --- |
| *Arachis duranensis* | 3 |
| *Cajanus cajan* | 3 |
| *Cicer arietinum* | 4 |
| *Glycine max* | 9 |
| *Medicago truncatula* | 5 |
| *Phaseolus vulgaris* | 4 |
| *Pisum sativum* | 4 |
| *Vigna angularis* | 5 |
| Average | 4.62 |
| Non-legume plants |  |
| *Arabidopsis thaliana* | 2 |
| *Hordeon vulgare* | 2 |
| *Populus trichocarpa* | 2 |
| *Ricinus communis* | 3 |
| *Solanum tuberosum* | 2 |
| *Sorghum bicolor* | 3 |
| *Vitis vinifera* | 2 |
| *Zea mays* | 2 |
| Average | 2.25 |

**Figure S1.**
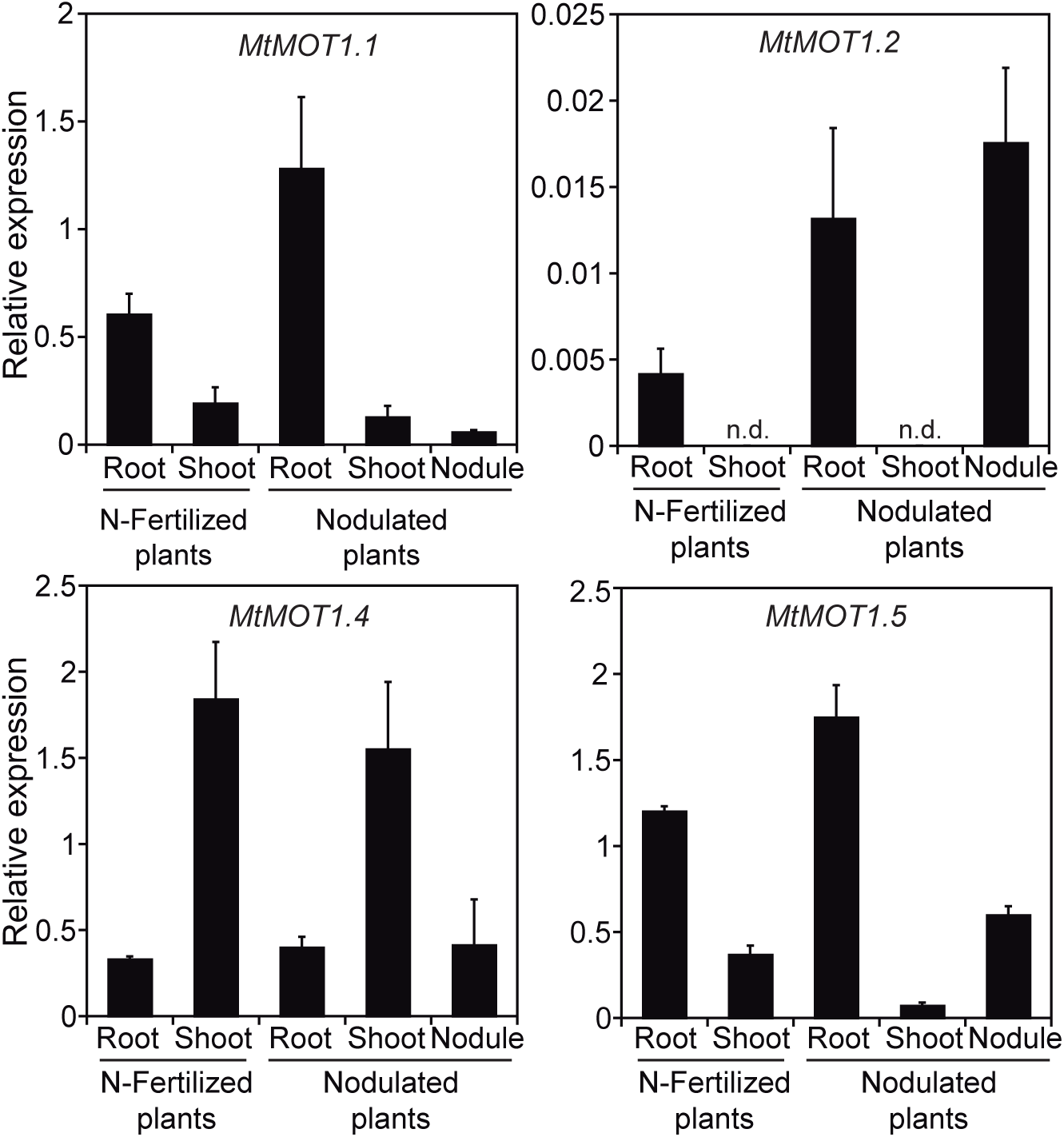
Determination of MOT1 gene family expression in nodulated and nitrogen-fertilized *M. truncatula* plants relative to the to the internal standard gene *Ubiquitin carboxyl-terminal hydrolase*. Data are the mean ± SD of two independent experiments with 4 pooled plants. n.d., non-detected.

**Figure S2.**
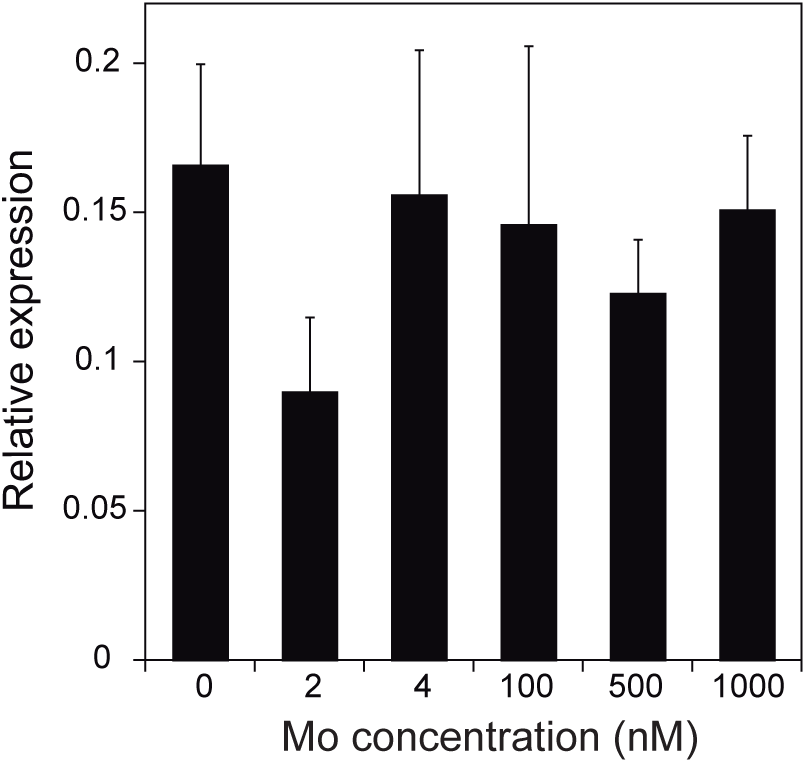
Determination of *MtMOT1.3* regulation by molybdate relative to the internal standard gene *Ubiquitin carboxyl-terminal hydrolase*. Data are the mean ± SD of three independent experiments with 4 pooled plants each.

**Figure S3.**
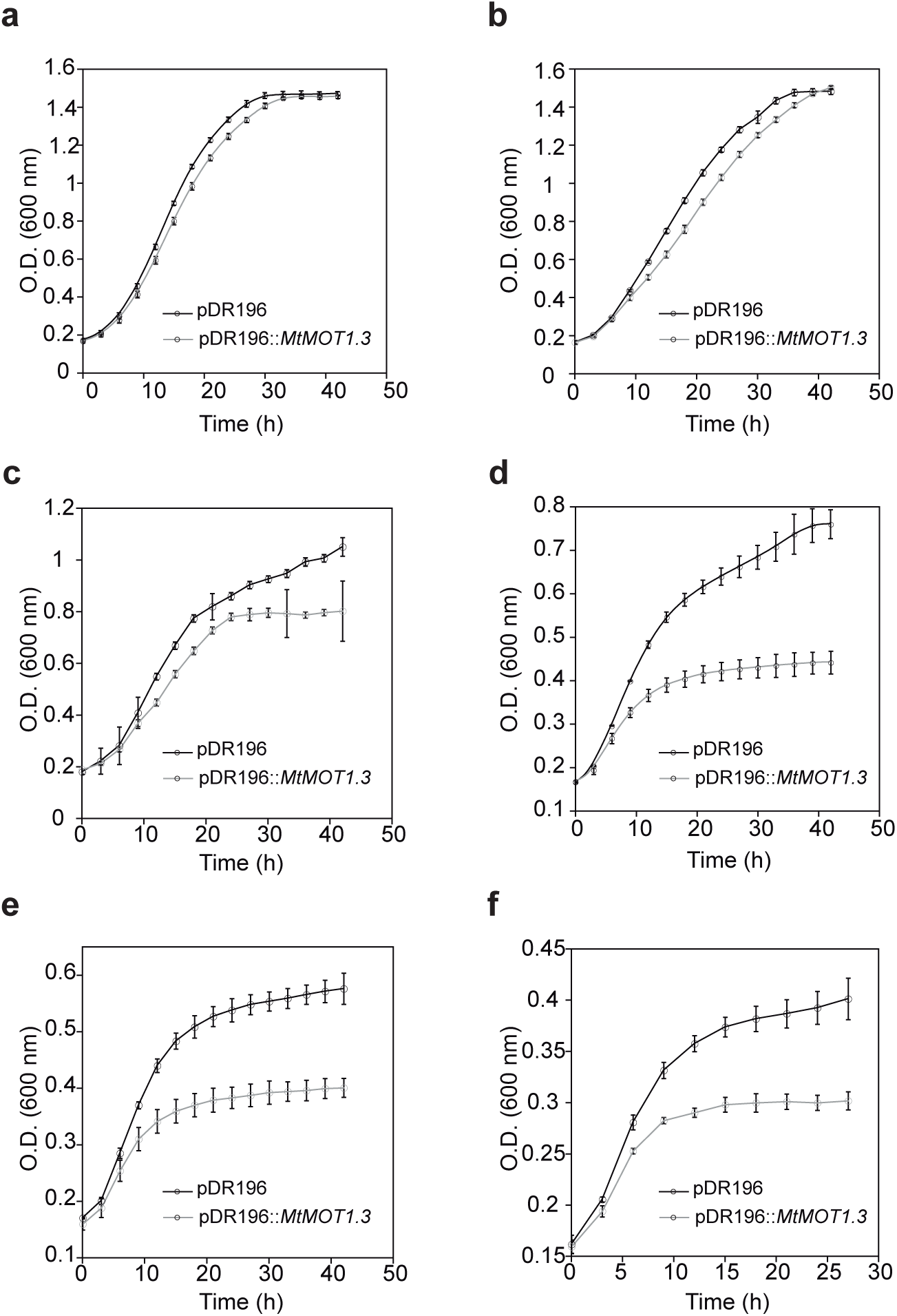
Growth of *S. cerevisiae* transformed with pDR196 or with pDR196 expressing MtMOT1.3 under different concentrations of sodium molybdate. (a) 0 µM added sodium molybdate. (b) 10 µM added sodium molybdate. (c) 30 µM added sodium molybdate.(d) 50 µM added sodium molybdate. (e) 100 µM added sodium molybdate. (f) 1000 µM added sodium molybdate. Data are the mean ± SD of two independent experiments.

**Figure S4.**
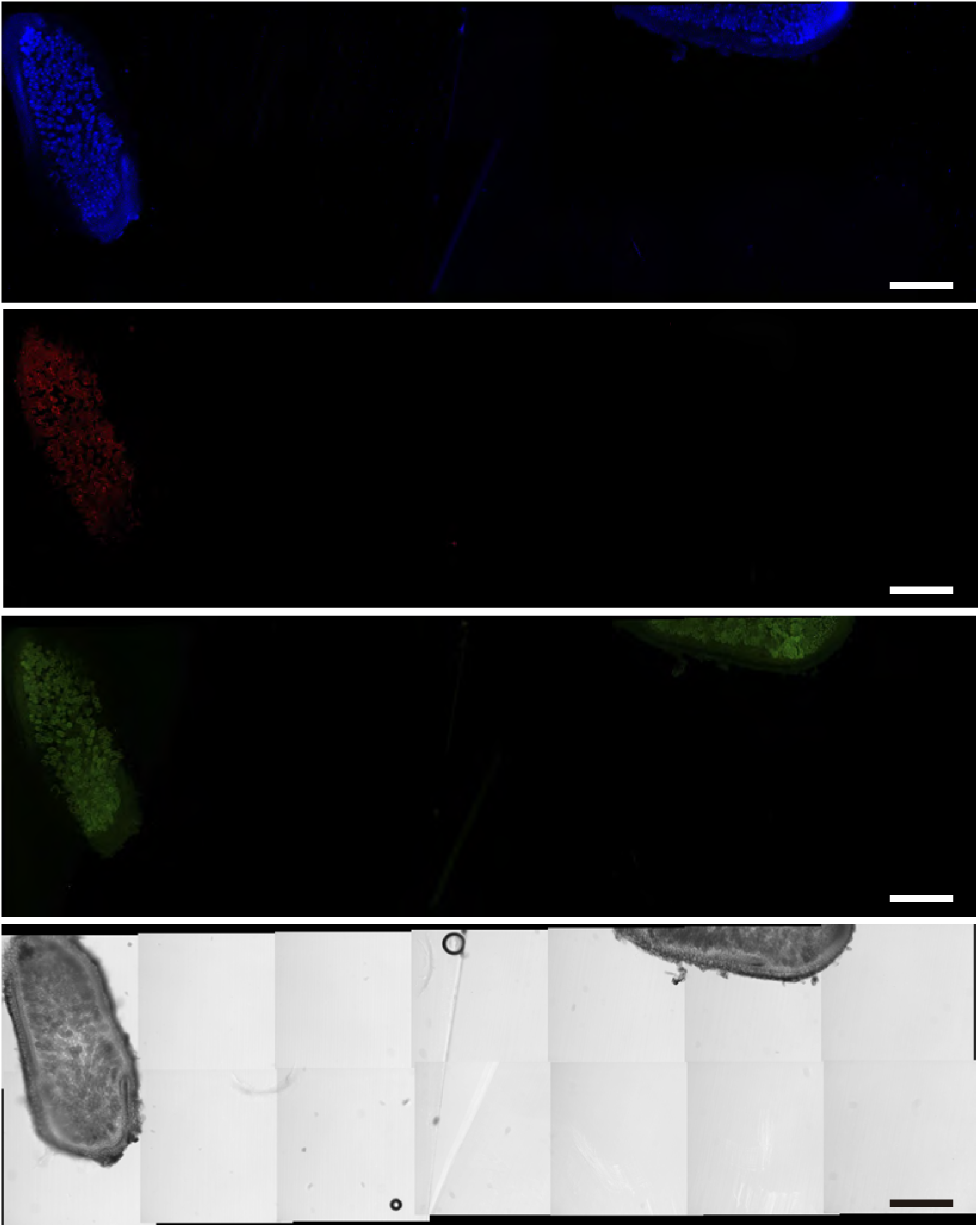
Autofluorescence control for Alexa594 emission. Immunolocalization of MtMOT1.3-HA in nodule sections from 28 dpi *M. truncatula* plants transformed with pGWB13::MtMOT1.3-HA. Left nodule was incubated with a mouse anti-HA antibody, while just PBS was used for the right one. The image is the result of a z-project of 22 optical sections combining a tilescan of 14 areas.

**Figure S5.**
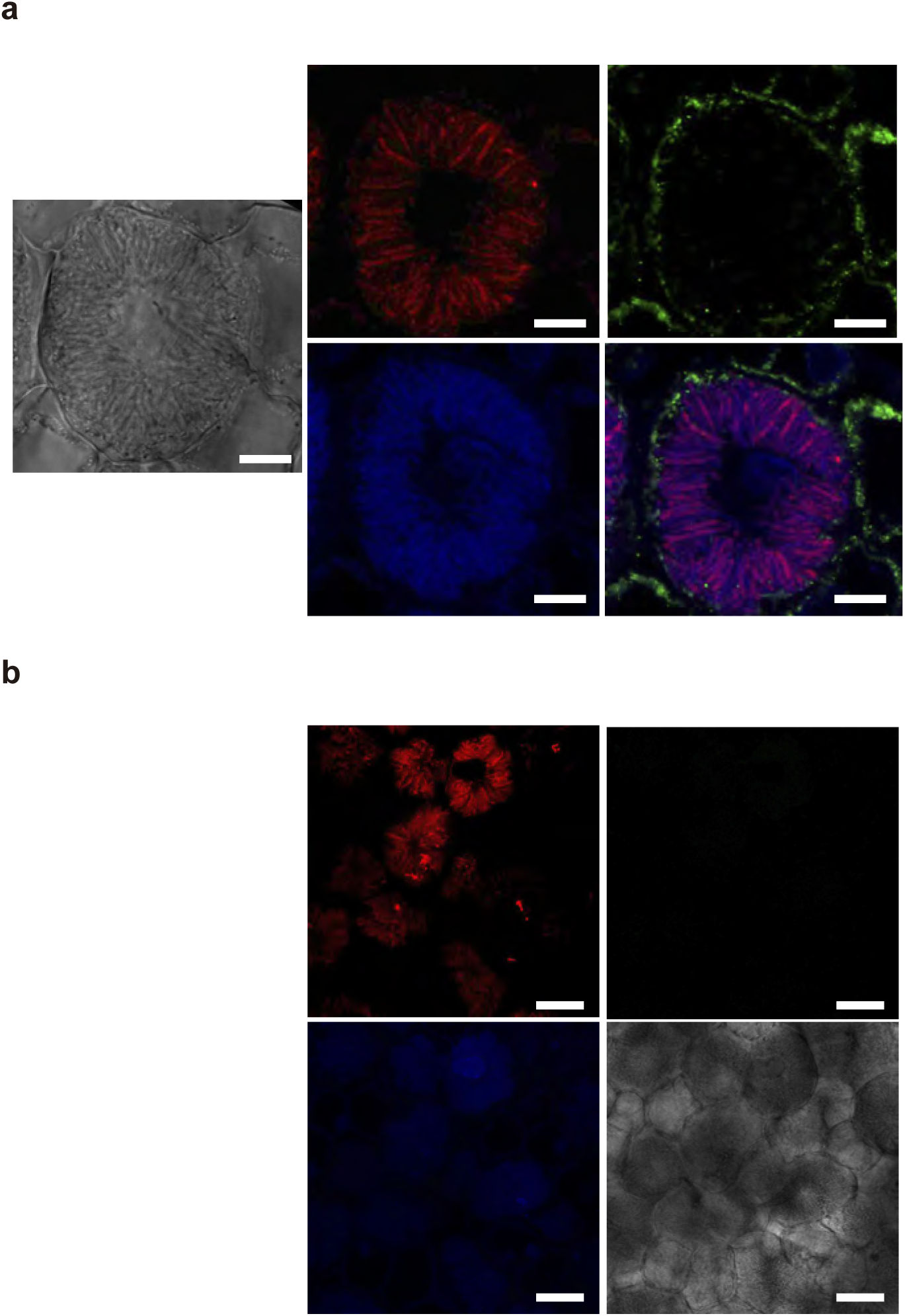
MtMOT1.3 is localized in the plasma membrane. Cross section of a 28 dpi *M. truncatula* nodule expressing MtMOT1.3-HA and infected with a S. *meliloti* strain constitutively expressing mCherry (red). DNA is stained with DAPI (blue). (a) Nodule section incubated with an anti-HA mouse antibody and a anti-mouse Alexa488-conjugated antibody (green). Left panel differential interface contrast image; top middle panel, S. meliloti distribution; lower middle panel, DNA distribution; top right panel MtMOT1.3-HA distribution; lower right panel overlay of DAPI, mCherry, and Alexa488 signals. Bars = 10 µm. (b) Nodule section prepared as in (a) but not adding the anti-HA mouse antibody. Top left panel, S. meliloti distribution; lower left panel, DNA distribution; top right panel distribution of Alexa488 signal; lower right panel differential interface contrast image. Bars = 20 µm.

**Figure S6.**
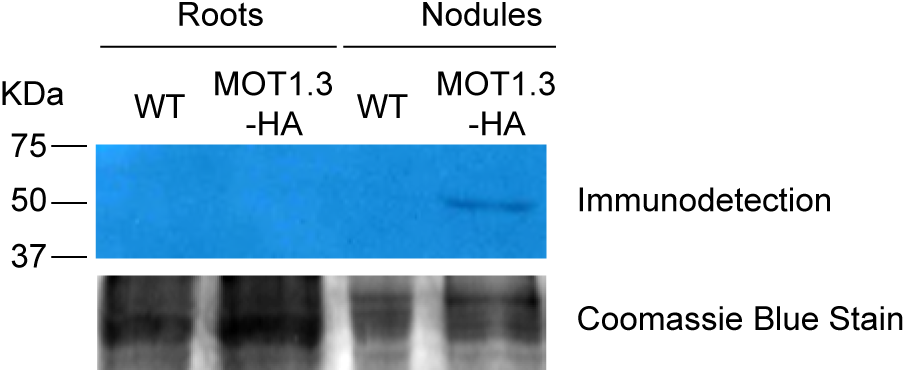
Immunolocalization of MtMOT1.3-HA in crude protein extracts from 28-dpi *M. truncatula* roots and nodules obtained from control and pGWB13::MtMOT1.3-HA transformed plants.

**Figure S7.**
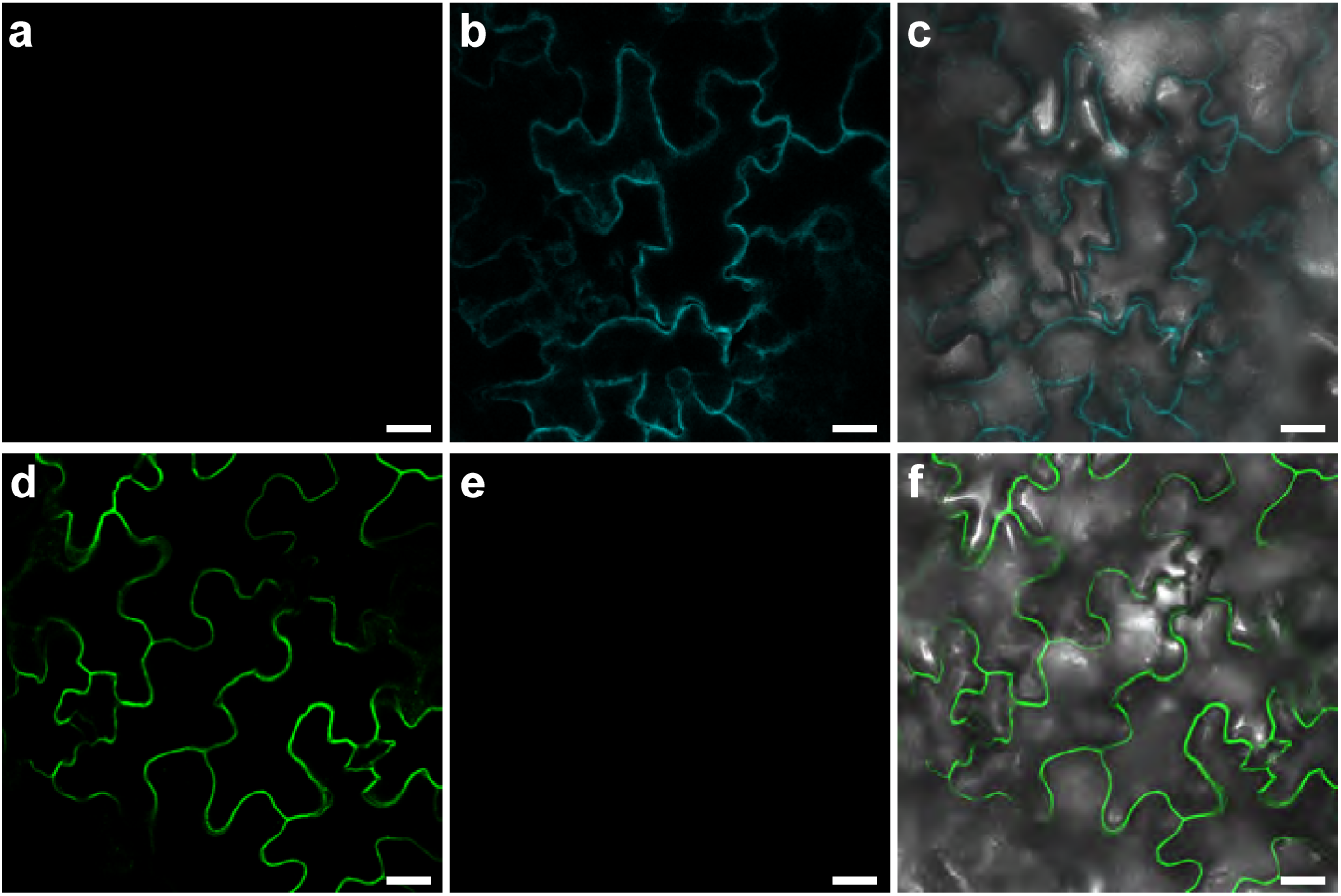
Transient expression of MtMOT1.3-GFP in *N. benthamiana* leaves. (a) to (c) Plasma membrane marker-CFP expression. (a) Plasma membrane marker-CFP signal in the green channel. (b) Plasma membrane marker-CFP in the cyan channel. (c) Overlay of green and cyan channels and differential interference contrast. Bars = 25 µm. (d) to (f) MtMOT1.3-GFP expression. (d) MtMOT1.3-GFP signal in the green channel. (e) MtMOT1.3-GFP signal in the cyan channel. (f) Overlay of green and cyan channels and differential interference contrast. Bars = 25 µm.

**Figure S8.**
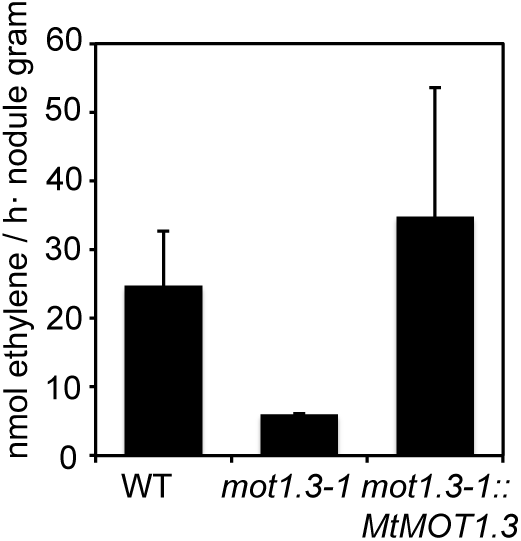
Nitrogenase activity in 28-dpi nodules. Acetylene reduction was measured in duplicate from two sets of four pooled plants. Data are the mean ± SD.

**Figure S9.**
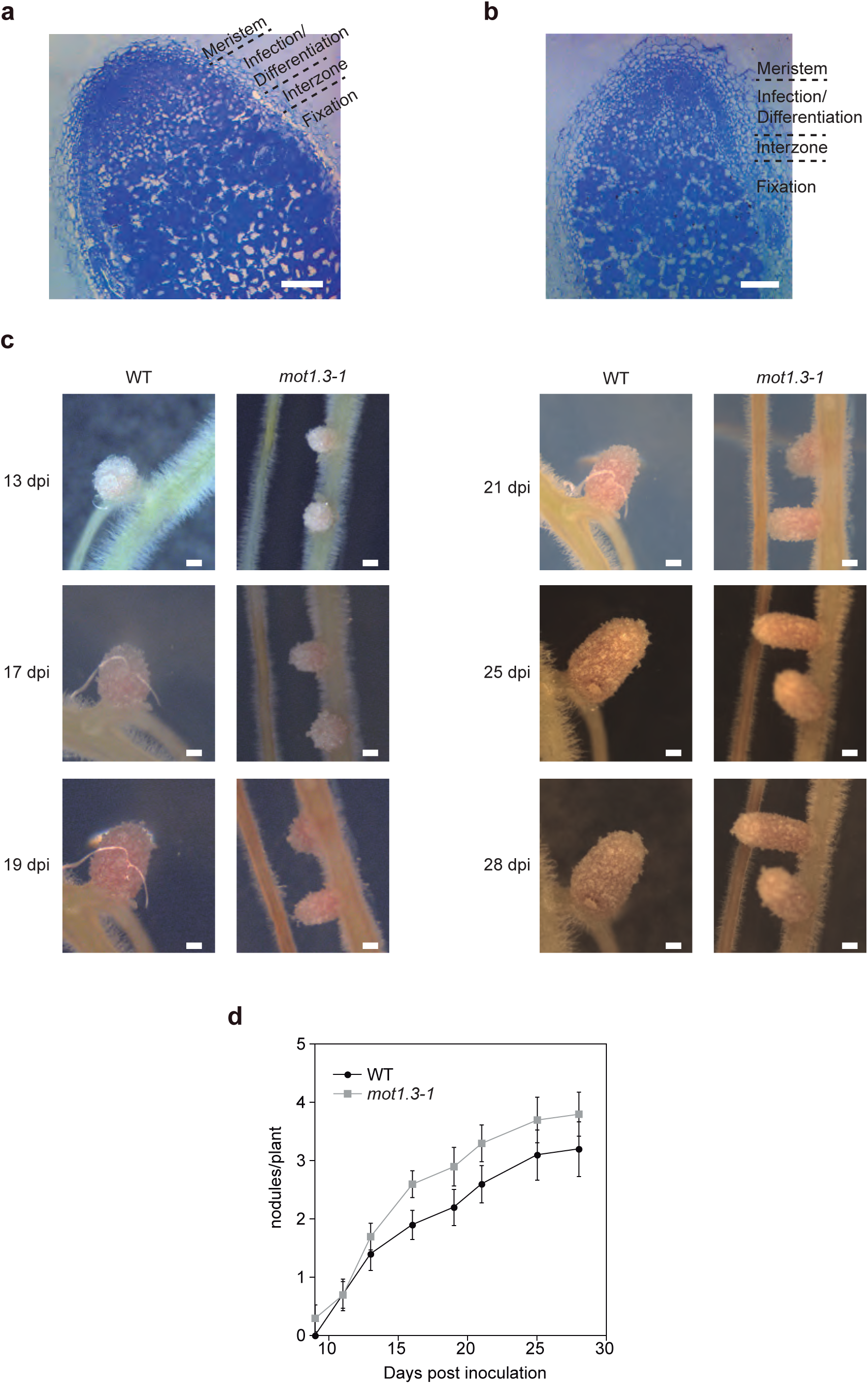
Nodule development in wild type and mot1.3-1 plants. Logitudinal sections of wild type (a) and mot1.3-1 (b) nodules stained with toluidine blue. Bars = 100 µm. (c) Nodule growth in wild type and mot1.3-1 plants. Bars = 100 µm (d) Nodulation kinetics in wild type and mot1-3 plants. Data are the mean ± SD from ten plants.

**Figure S10.**
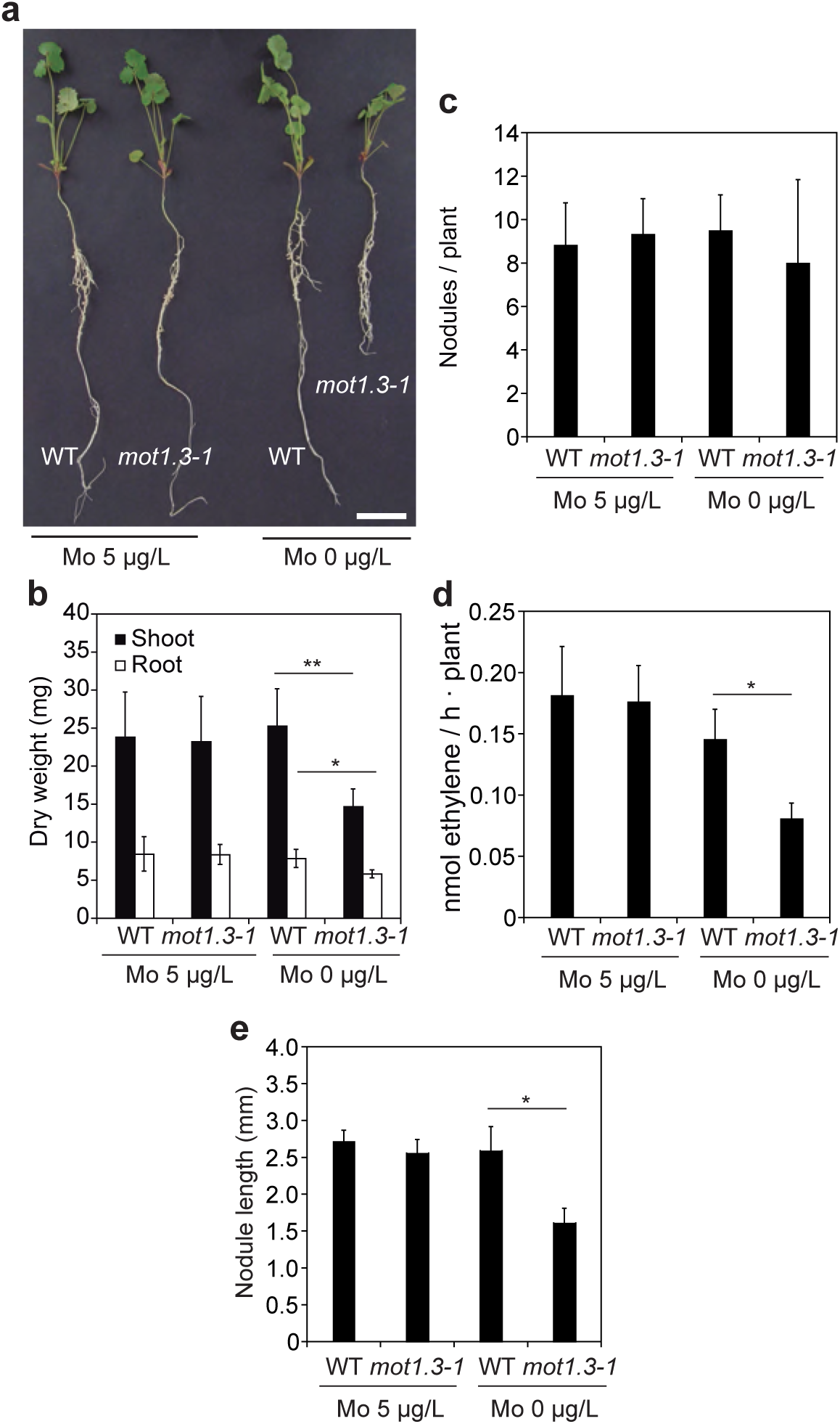
*mot1.3-1* mutant aberrant phenotype is restored upon supplementation with molybdate. (a) Growth of representative plants of wild type and *mot1.3-1* mutant. Bar = 3 cm. (b) Dry weight of shoots and roots. Data are the mean ± SD of at least 6 independent plants. Asterisk indicates significant differences: *p < 0.05, **p < 0.01. (c) Number of nodules per plant. Data are the mean ± SD of at least 6 independent plants. (d) Nitrogenase activity in 28-dpi nodules. Acetylene reduction was measured in duplicate from two sets of four pooled plants. Data are the mean ± SD. Asterisk indicates significant differences: *p < 0.05. (e) Nodule length in 28-dpi nodules. Data are the mean ± SD of 15 nodules from from 6 plants. Asteriks indicates significant differences: *p < 0.05.

**Figure S11.**
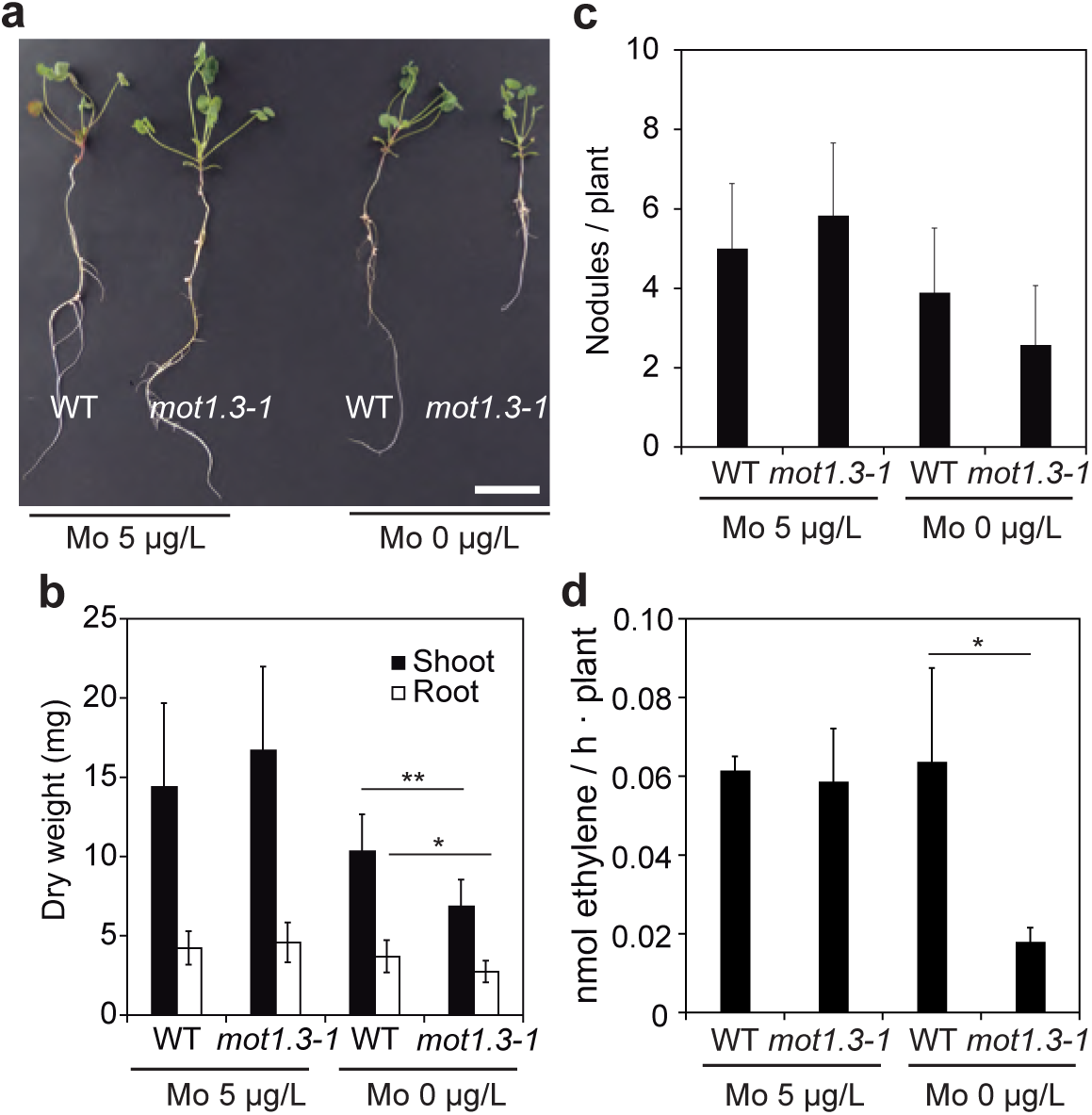
*mot1.3-1* plants show reduced nitrogenase activity also in hydroponics. (a) Growth of representative plants of wild type and mot1.3-1 mutant. Bar = 3 cm. (b) Dry weight of shoots and roots. Data are the mean ± SD of at least 8 independent plants. Asterisk indicates signifi-cant differences: *p < 0.05, **p < 0.01. (c) Number of nodules per plant. Data are the mean ± SD of at least 7 independent plants. (d) Nitrogenase activity in 28-dpi nodules. Acetylene reduction was measured in duplicate from two sets of four pooled plants. Data are the mean ± SD. Asterisk indicates significant differences: *p < 0.05

**Figure S12.**
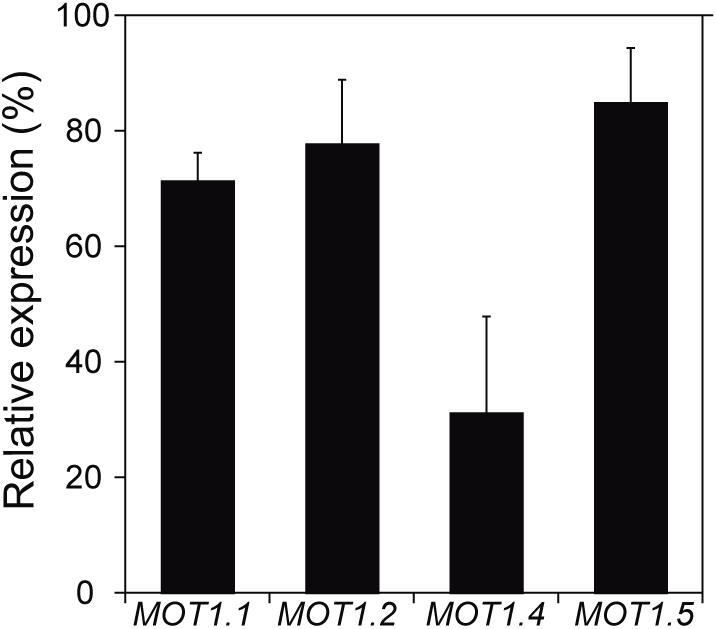
Expression of *MtMOT1.1, MtMOT1.2, MtMOT1.4,* and *MtMOT1.5* in *mot1.3-1* nodules relative to the internal standard gene *Ubiquitin carboxyl-terminal hydrolase* and relativised to the expresion value in wild type plants (100 %). Data are the mean ± SD of three independent experiments with 4 pooled plants each.

